# Placeholder nucleosomes underlie germline-to-embryo DNA methylation reprogramming

**DOI:** 10.1101/204792

**Authors:** PJ Murphy, SF Wu, CR James, CL Wike, BR Cairns

**Affiliations:** Howard Hughes Medical Institute, Department of Oncological Sciences and Huntsman Cancer Institute, University of Utah School of Medicine, Salt Lake City, UT 84112, USA

**Keywords:** DNA methylation, zebrafish, H2A.Z, germline-to-embryo transition

## Abstract

The fate and function of epigenetic marks during the germline-to-embryo transition is a key issue in developmental biology, with relevance to stem cell programming and trans-generational inheritance. In zebrafish, DNA methylation (DNAme) patterns are programmed in transcriptionally-quiescent cleavage embryos; remarkably, paternally-inherited patterns are maintained, whereas maternal patterns are reprogrammed to match the paternal. Here we provide the mechanism, by demonstrating that ‘Placeholder’ nucleosomes, containing histone H2A variant H2A.Z(FV) and H3K4me1, occupy virtually all regions lacking DNAme in both sperm and cleavage embryos, and resides at promoters encoding housekeeping and early embryonic transcription factors. Upon genome-wide transcriptional onset, genes with Placeholder become either active (H3K4me3) or silent (H3K4me3/K27me3). Importantly, functional perturbation causing Placeholder loss confers DNAme accumulation, whereas acquisition/expansion of Placeholder confers DNA hypomethylation and improper gene activation. Thus, during transcriptionally-quiescent gametic and embryonic stages, an H2A.Z(FV)/H3K4me1-containing Placeholder nucleosome deters DNAme, poising parental genes for either gene-specific activation or facultative repression.

**Highlights: four, 85 characters each:**

- Placeholder nucleosomes bear H3K4me and the histone variant H2A.Z(FV)
- Placeholders occupy all DNA hypomethylated loci in zebrafish sperm and early embryos
- At ZGA, Placeholders resolve into either active or poised (bivalent) nucleosomes
- Placeholder localization excludes DNAme, and regulates embryonic transcription

## Introduction

A major question in early embryos is how chromatin packaging of the parental genomes helps ensure proper gene regulation during and after genome-wide transcriptional activation. Chromatin packaging differs greatly between mature sperm and eggs, yet these differences are largely harmonized (imprinted loci excluded) via reprogramming in the early vertebrate embryo (Akkers et al., 2009; Dahl et al., 2016; Hammoud et al., 2009; Liu et al., 2016; Vastenhouw et al., 2010; Wu et al., 2011; Zhang et al., 2016). The zebrafish (*Danio rerio*) model system offers an opportunity to study many aspects of this process in detail due to the abundant availability of sperm, eggs and embryos - and key properties of zebrafish cleavage-stage embryos. Unlike mammals, prior to initiating genome-wide zygotic genome transcriptional activation (ZGA), zebrafish undergo 10 rapid cell divisions during cleavage phase resulting in a ~4000-cell largely pluripotent embryo at 4 hours post-fertilization (4hpf), undergoing active transcription (Lee et al., 2013).

Prior work revealed several key features regarding the reprogramming of parental epigenomes during the gamete-to-embryo transition in zebrafish. Both the paternal and maternal zebrafish genomes are packaged solely in histones (not protamines) in mature sperm and eggs (Wu et al., 2011; Zhang et al., 2016), and similar to mammals, the zebrafish sperm genome appears ‘poised’ for the future activation of particular gene categories. For example, sperm lack DNA methylation (DNAme) at the promoters of housekeeping genes (H3K4me3-marked), and also at bivalent genes (H3K4me3 and H3K27me3-marked (Bernstein et al., 2006)) that encode transcription factors (TFs) for guiding somatic development (e.g. *hox*-family, *gata-* family factors), pluripotency factors (e.g. *nanog* and *pou5f3* (the *zebrafish* OCT4 *ortholog*)) and genes for germline identity/function (e.g. *dazl*, *piwil1*) (Wu et al., 2011, Potok et al., 2013). Thus, the paternally-inherited DNAme patterns of zebrafish sperm are compatible with achieving pluripotency and with the derivation of germline. Likewise, zebrafish oocytes lack DNAme at housekeeping genes and certain developmental genes; however, they possess clear DNAme at several hundred loci important for development, pluripotency, and germline identity/function (e.g. many *hox*-family genes, *nanog*, *dazl*, *piwil1*). Whereas the paternal DNAme patterns are virtually identical to those in pluripotent embryos at 4hpf, the maternal genome is instead gradually reprogrammed to the paternal pattern during cleavage (Jiang et al., 2013; Potok et al., 2013). Presently, the mechanisms that enable maintenance of paternal DNAme and reprogramming of maternal DNAme in zebrafish embryos are unknown, and may inform similar issues in mammals.

In other systems, achieving DNA hypomethylation (hypoDNAme) occurs via a host of mechanisms; however, many of these cannot account for DNAme reprograming in early zebrafish embryos. For example, H3K4-methylation and/or H3K27me3 antagonize DNAme (Brinkman et al., 2012; Murphy et al., 2013; Ooi et al., 2007; Zhang et al., 2010), yet cleavage-stage zebrafish embryos display strong focal hypoDNAme in the absence of high H3K4me3 and/or H3K27me3, which are highly depleted during cleavage (Lindeman et al., 2010; Vastenhouw et al., 2010; Zhang et al., 2014). Bivalent marks are also diminished in early mouse embryos (Dahl et al., 2016; Liu et al., 2016; Zhang et al., 2016); thus, alternative factors or modifications may be needed to maintain focal chromatin information in mammalian embryos. Moreover, the use of TET-family enzymes and 5hmC (Hackett et al., 2013; Huang et al., 2014; Tahiliani et al., 2009) is also excluded in cleavage-stage zebrafish, as they are absent this phase (Potok et al., 2013) and instead utilized at/after gastrulation (Bogdanović et al., 2016). Here, we provide a mechanism for establishing and maintaining focal DNA hypoDNAme during cleavage, involving a nucleosome bearing the histone H2A variant H2A.Z(FV) and H3K4me1.

The histone H2A variant H2A.Z (termed H2AFV in zebrafish) is similar to canonical H2A, but differs along its amino terminal ‘tail’ and at key residues in the globular domain and C-terminus. Precise genomic incorporation of H2A.Z occurs by the nucleosome remodeler SRCAP (Snf2-Related CREBBP Activator Protein) (Kobor et al., 2004; Krogan et al., 2003; Wang et al., 2016). H2A.Z installation is antagonized by the recently-identified chaperone Anp32e, which conducts H2A.Z removal (Mao et al., 2014; Obri et al., 2014). In vertebrates, H2A.Z-containing nucleosomes reside at most gene promoters, including developmentally-regulated genes (Barski et al., 2007; Hu et al., 2013; Ku et al., 2012), and H2A.Z has roles in differentiation, nucleosome stability, transcriptional regulation, OCT4 binding and Polycomb-complex recruitment (Chen et al., 2014; Creyghton et al., 2008; Hu et al., 2013; Jin et al., 2009; Ku et al., 2012). However, the role of H2A.Z in developmental epigenetic reprogramming during the germline-to-embryo transition remains largely unexplored. The relationship between H2A.Z and DNAme is complex, but often antagonistic. In plant studies (*Arabidopsis thaliana*) DNA methyltransferase (DNMT) mutants display decreased DNAme and increased H2A.Z occupancy, and mutants for the H2A.Z-specific nucleosome remodeler *swr1* (an SRCAP ortholog) have decreased H2A.Z incorporation and increased DNAme over gene bodies (Zilberman et al., 2008). Furthermore, in mammalian cancer cells, H2A.Z antagonizes DNAme during MYC-mediated mouse B-cell transformation (Conerly et al., 2010), which causes redistribution of H2A.Z from promoters to gene bodies along with increased DNAme at promoters and decreased DNAme over gene bodies.

Here, we utilized zebrafish to address a key question in vertebrate germline-to-embryo epigenetics: How are paternal DNAme patterns maintained, and maternal DNAme patterns reprogrammed during a transcriptionally-quiescent cleavage phase – when bivalent chromatin is largely lacking? Taken together, our work provides evidence for ‘Placeholder’ nucleosomes – which consists of zebrafish H2A.Z (H2AFV) and H3K4me1 – to ‘hold in place’ hypoDNAme regions, which then resolve at ZGA into either hypoDNAme active or poised chromatin. Through a series of gain-of-function and loss-of-function approaches we demonstrate the molecular utility of Placeholder in programming *in vivo* DNAme patterns, and in the regulation of transcriptional activation during early zebrafish development.

## Results

### Specific chromatin features mark DNA hypomethylated regions in sperm

DNA hypomethylated regions in sperm and early embryos are striking in their sharp boundaries (Sup Figure 1a). To account for this precision of focal DNAme reprogramming during zebrafish cleavage stage we reasoned that a ‘Placeholder’ type of nucleosome/chromatin – bearing particular histone variants and/or modifications – might exist to antagonize DNAme in both sperm and transcriptionally-quiescent embryos (Figure 1a). We identified candidate ‘Placeholder’ marks/features by genome-wide chromatin immunoprecipitation followed by sequencing (ChIP-Seq) to profile known promoter (H3K4me3, H3K27me3, H3K14ac and zebrafish H2A.Z/FV) and enhancer features (H3K4me1 and H3K27ac) in zebrafish sperm. These marks showed the expected profiles at TSS and gene body regions (Figure 1b & Sup Figure 1a). K-means clustering of genome-wide data revealed low DNAme overlapping with high levels of these marks (Figure 1c – cluster 2, & Sup Figure 1b) (DNAme data from Potok et al., 2013). However, DNAme appeared compatible with robust levels of H3K4me1, and low-moderate H3K4me3 (cluster 1), indicating their insufficiency (alone, at those levels) for deterring DNAme in sperm. In counter distinction, virtually all regions with high H2AFV, and H3K14ac lacked DNAme (cluster 2, & Sup Figure 1c).

**Figure 1:**
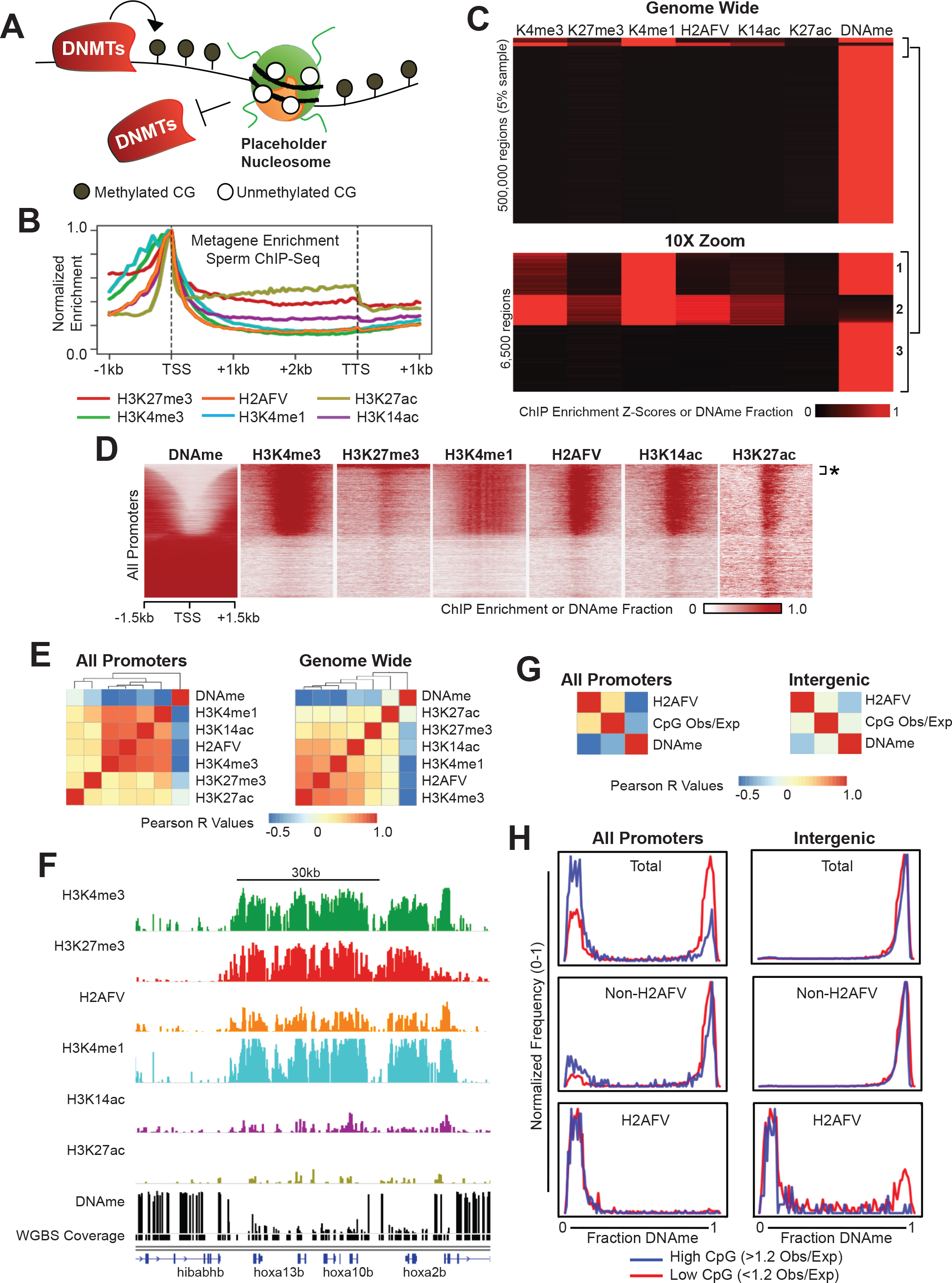
Identification of Placeholder features in zebrafish sperm chromatin. A) A model for ‘Placeholder’ nucleosome blocking DNA methyltransferases (DNMTs) where it resides to facilitate hypoDNAme. B) Metagene analysis of normalized ChIP-Seq enrichment. Epigenetic features are enriched at the transcription start site (TSS) and depleted within the gene body. C) K-means clustering of epigenetic features in sperm. Top - genome-wide; bottom - 10X zoom for higher resolution. Z-scores are displayed for ChIP-Seq data and fraction methylation for DNAme. All regions are 200bp. D) H3K4me3, H3K4me1, H2AFV, and H3K14ac mark hypoDNAme promoters in sperm. Heatmaps of Z-scores are displayed for ChIP-Seq data and fraction methylation for DNAme. * = bivalent loci. E) H3K4me3, H3K4me1, H2AVF, and H3K14ac are correlated with one another, and anti-correlated with DNAme in sperm. Clustering of pairwise Pearson correlation R values are displayed as a heatmap. F) A genome browser snapshot of the *hoxA* locus. Log10FE (0-1) values for ChIP-Seq data and DNAme fraction (0-1). G) H2AFV is correlated with CpG density only at promoters. Pairwise correlation analysis as in panel-E, specifically focused on promoters (left) and intergenic loci (right). H) H2AFV marks hypoDNAme regions regardless of CpG density. Histograms indicate frequency after parsing regions for H2AFV, DNAme level, and CpG density. Left - promoters, and right - intergenic regions.

Similarly, the majority (55%) of sperm promoters lacked DNAme, and were generally marked with one or more of these modifications, most prominently high H2AFV or H3K4me1/3 (Figure 1c, cluster 2; Figure 2d). Notably, promoters with high H3K27me3 and H3K4me3 (bivalent) also lacked DNAme, but represented only a small fraction (~5%) of total promoters (Figure 2d, asterisk). Pearson correlational analyses revealed H3K4me3, H2AFV and H3K4me1 as the features most strongly anti-correlated with DNAme both genome-wide and at promoters (Figure 1e) – exemplified by key developmental genes (e.g. *hoxa* cluster, Figure 1f). DNAme is anti-correlated with CpG density in mammals and zebrafish (Potok et al., 2013), and although H2AFV occupancy correlated with CpG density at promoters (Figure 1g, left panel), somewhat unexpectedly, they are uncorrelated outside of promoters (Figure 1g, right panel). To further investigate, we parsed the genome into promoters vs. intergenic loci and examined H2AFV occupancy, CpG density and DNAme (Figure 1h). Again, promoters with high CpG density were H2AFV occupied and lacked DNAme (Figure 1h - top left). In contrast, at intergenic regions high DNAme is observed regardless of CpG density (Figure 1h - top right). Interestingly, regions that lack H2AFV bear high DNAme (Figure 1h - middle), whereas those marked by H2AFV lack DNAme, regardless of promoter/intergenic context or CpG density (Figure 1h - bottom). These results demonstrate that H2AFV occupancy in sperm is more strongly anti-correlated with, and predictive of, DNAme state than CpG richness. Taken together, H2AFV is strikingly anti-correlated with DNAme, and is often found with H3K4me1 (exemplified in Sup Figure 1a).

**Figure 2:**
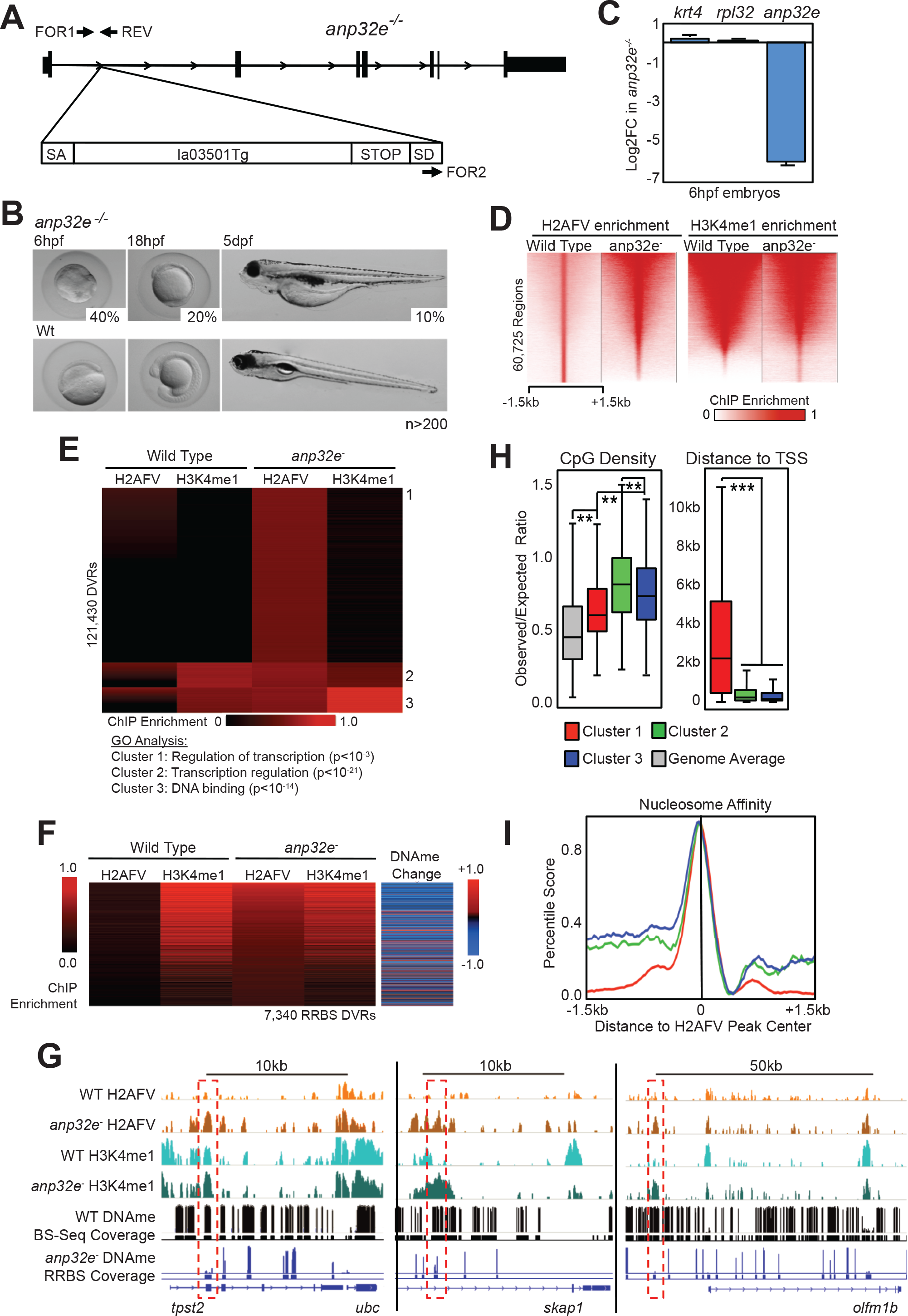
A null mutation for *anp32e* causes increased genomic H2AFV and decreased DNAme in sperm. A) Scheme for the *anp32e* mutant allele. A retroviral gene trap cassette (la013501Tg) within the first intron of the *anp32e* gene causes premature translation termination. PCR primers for genotyping are indicated. B) *anp32e*^−/−^ homozygotes display a variety of phenotypes. Lethality occurs at the displayed stages. The remaining 30% are viable and fertile. C) *anp32e* mRNA in the *anp32e* mutants (log fold change). An early zygotically-transcribed gene (*krt4*) and a housekeeping gene (*rpl32*) are unaffected. D) H2AFV enrichment regions become more broad in *anp32e*^-^ sperm. Heatmaps for H2AFV and H3K4me1 are displayed at all WT sperm H2AFV peaks. E) *anp32e*^-^ sperm display ectopic H2AFV localization. H2AFV and H3K4me1 enrichment is displayed as a heatmap in WT and mutant sperm. Differential Variant Regions (DVRs) were analyzed and segregate into 3 clusters by K-means clustering. Results of GO analysis for affected gene categories is shown below. F) DNAme decreases at the majority of DVRs. Displayed regions are only those with sufficient read-depth from RRBS-Seq assays (min. 8 reads and 3 CpG per 50bp). Key on right side of panel indicates changing DNAme in mutant sperm compared to WT. G) A gallery of browser snapshots is displayed for WT and *anp32e*^-^ sperm. Regions with sufficient coverage for WGBS DNAme data are indicated with black bars underneath, or for RRBS with blue bars underneath. Ectopic H2AFV accumulation is highlighted with red dashed boxes. Log10FE data from 0-1 for ChIP-Seq, and DNAme fraction from 0-1. H) Analysis of CpG density (left) and neighboring gene proximity distance (right) for clusters identified in panel E. T-tests (see methods): ** = p< 0.01, *** = p < 0.001. I) Analysis of predicted nucleosome affinity at DVRs. Regions from panel E were grouped by cluster and colored as in panel G.

### Acquisition of H2AFV confers loss of DNAme in sperm

To test whether H2AFV functionally regulates DNAme patterns we performed loss-of-function assays. Here, the ubiquitous expression and separate paralogues of *h2afv* genes in the zebrafish genome deterred a genetic knockout approach. Therefore, we disrupted the singlecopy gene, *Srcap*, which encodes the conserved catalytic subunit of the chromatin remodeler SRCAP complex (Kobor et al., 2004; Krogan et al., 2003; Wang et al., 2016). Our TALEN-mediated disruption (Dahlem et al., 2012) of *srcap* was successful; however, homozygous null animals failed to survive to sexual maturity (phenotypes to be described later). Transplantation of *srcap*^−/−^ germline stem cells into wild type (WT) surrogate hosts yielded solely males that lacked sperm, which typically occurs when donor germline stem cells die, likely due to *srcap* performing roles essential for cell survival. Therefore, as an alternative strategy we isolated zebrafish mutants lacking Anp32e, the histone chaperone known to remove H2A.Z in mice (Obri et al., 2014).

As zebrafish contain a close ortholog of *Anp32e*, zebrafish *anp32e* null mutants (if viable) would offer the potential to study the impact of H2A.Z mislocalization in maternal zygotic Anp32e deficient vertebrate embryos. We obtained the available heterozygous *anp32e* null mutation (transgenic la013501 insertion; Figure 2a and Sup Figure 2a) (Varshney et al., 2013). Crossing generated homozygous animals that displayed phenotypic variation - including lethality, which occurred along developmental stages: shield stage (~40%), early somite (~20%), or 5 days post fertilization (dpf; ~10%) (Figure 2b). However, ~30% of the embryos survived to adulthood, genotyped as *anp32e*^−/−^ lacked detectable *anp32e* RNA at shield stage (unlike WT) (Figure 2c), and were fertile – providing a source of *anp32e*^-^ sperm and oocytes.

As H3K4me1 and H2AFV were largely coincident at hypoDNAme regions, we investigated their genomic localization upon Anp32e loss. First, we observed a clear broadening of H2AFV peaks in *anp32e*^-^ mutant sperm relative to WT sperm (Figure 2d). Here, mutant-specific enrichment for H2AFV occurred largely at intergenic regions, and overall enrichment localization remained constant for H3K4me1 (Sup Figure 2b). We identified over 120,000 *de novo* ectopic H2AFV regions in *anp32e*^-^ mutant sperm - defined as differential variant regions (DVRs) (see Methods). Relationships between H2AFV and H3K4me1 at these sites were revealed through K-means clustering, which yielded three clusters (Figure 2e): cluster 1 loci contain H2AFV but lack H3K4me1, loci within clusters 2 and 3 bear moderate-high H2AFV and H3K4me1, and cluster 3 bears the highest H3K4me1. Thus, a portion of the sites that acquire H2AFV also acquire H3K4me1. Notably, examination of gene categories (via GO analysis) that acquire ectopic H2AFV yielded enrichment at genes encoding DNA-binding transcriptional regulators (Figure 2e, bottom).

Next, we determined the impact of increased H2AFV on DNAme by performing Reduced Representation BiSulfite Sequencing (RRBS-Seq). Strikingly, the vast majority (71%) of regions that acquired ectopic H2AFV lost DNAme (Figure 2f), displaying a median reduction in fraction DNAme of 13%. Cluster 1, which presented the largest increase in H2AFV, exhibited the greatest decrease in DNAme (median level lowered from 59% to 32%). Examples of affected loci are provided in Figure 2g (note: affected regions that meet RRBS coverage thresholds are highlighted, see legend). Having revealed a strong effect on DNAme, we then explored correlations with CpG content, and found that the H2AFV/H3K4me1 clusters from Figure 2e differed in CpG content: cluster 1 regions were less CpG rich, and located distal to gene promoters, whereas clusters 2 and 3 were CpG rich and promoter proximal (Figure 2h). Thus, CpG-dense promoter-proximal regions that were marked by H3K4me1 and lacked H2AFV in WT, gained H2AFV in *anp32e* mutant sperm; whereas at CpG-poor distal intergenic regions that gained H2AFV in *anp32e* mutant sperm lacked H3K4me1 in WT sperm. CpG density and GC richness can affect nucleosome stability, and sequences that stabilize or destabilize nucleosomes can be predicted *in silico* (Kaplan et al., 2009). Remarkably, we observed a high nucleosome affinity score at the center of ectopic H2AVF peaks in all three clusters (Figure 2i). Taken together, Anp32e loss causes ectopic acquisition of H2AFV in sperm at regions of high nucleosome affinity and these regions have deeply diminished DNAme compared to WT.

Similar approaches with oocytes (either WT or *anp32e*^−^) were prevented, as H2AFV levels were low, and we were unable to enrich for H2AFV in oocytes by ChIP (data not shown). The diffuse pattern of H2AFV staining in oocytes (lacking clear chromatin association) (Pauls et al., 2001) and high levels of maternal Anp32e (data not shown) suggests that oocytes may lack appreciable chromatin-bound Placeholder. Instead, we find focal hypoDNAme in oocytes co-incident with high H3K4me3, present at housekeeping gene promoters, and absent at loci for developmental transcription factors (TFs) (Sup Figure 3a&b) (Zhang et al., 2016). Thus, oocytes may utilize focal H3K4me3 to achieve focal hypoDNAme – and reserve the use of Placeholder for maternal genome DNAme reprogramming during cleavage, explored below.

### Locations of H2AFV in sperm are largely maintained in embryos

Having demonstrated that H2AFV restricts DNAme in sperm, we next examined whether H2AFV locations were maintained in embryos to retain paternal DNAme patterns during cleavage stage. To explore this, we performed genome-wide ChIP-Seq in embryos for H2AFV, H3K4me1, and H3K14ac, as well as many additional modifications (Sup Figure 3c). ChIP-Seq was performed at two time points: well prior to ZGA (2.5 hpf), or just after ZGA (4hpf). Notably, we attempted ChIP-Seq for H3K4me3 and H3K27me3 in preZGA samples but, similar to observations by others (Vastenhouw et al., 2010; Zhang et al., 2014), we detected no enrichment for these marks. We also compared our profiles to published postZGA profiles of H3K4me3, H3K27me3, H3K4me1, and H3K27ac (Bogdanovic et al., 2012; Zhang et al., 2014). Interestingly, locations with H2AFV and H3K4me1 (but not H3K14ac) in sperm correlated well with those locations in embryos, and similar to our sperm results, embryonic H2AFV and H3K4me1 were strongly anti-correlated with embryonic DNAme (Figure 3a). Furthermore, immunoprecipitation of H2AFV (primarily mononucleosomes, via MNase treatment) from preZGA embryo chromatin (2.5hpf) co-precipitated H3K4me1, supporting the co-incidence of both H3K4me1 and H2AFV on individual nucleosomes in embryos (Sup Figure 3d).

**Figure 3:**
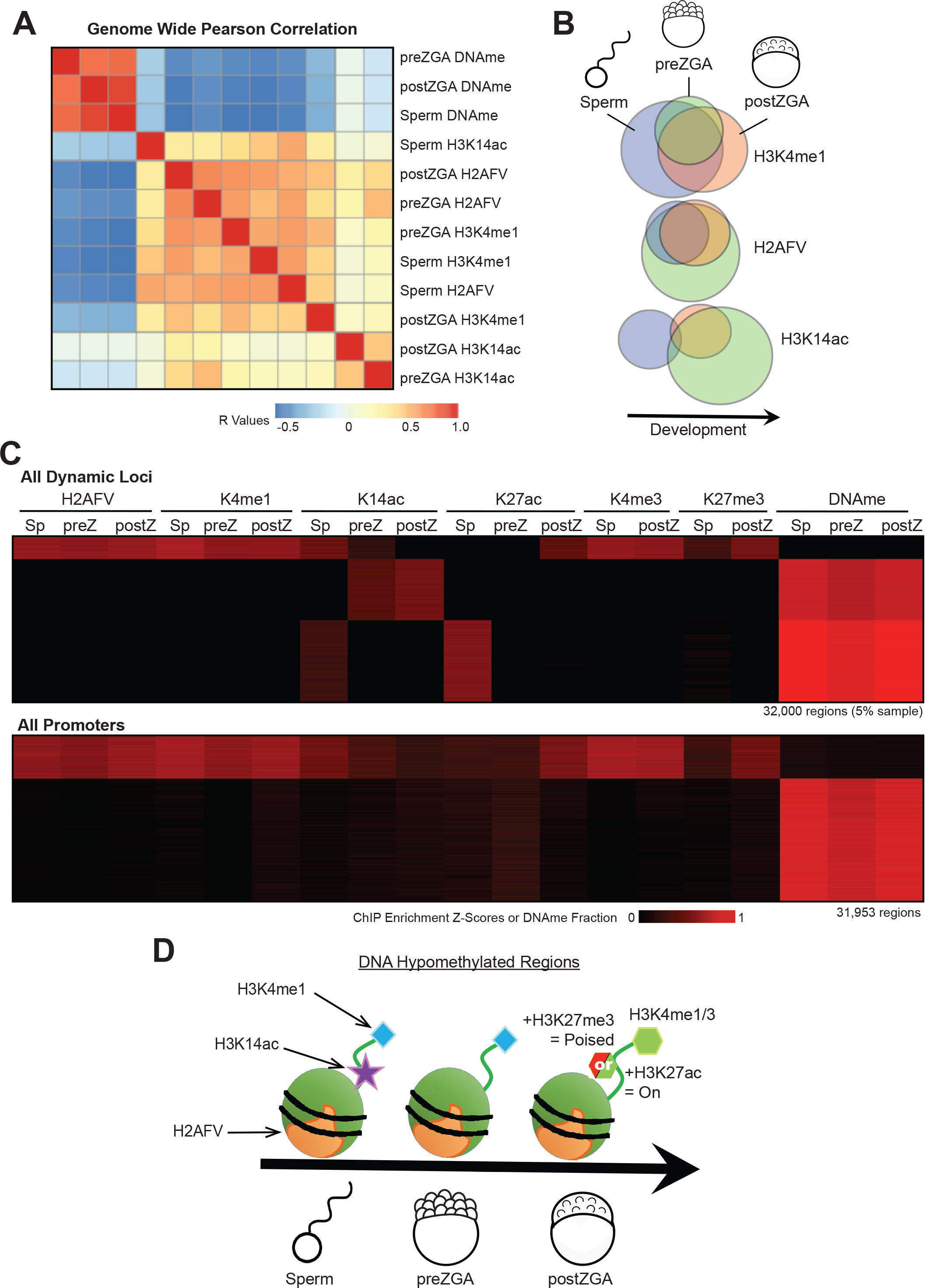
Maintenance of H2AFV and H3K4me1 at DNA hypomethylated regions in zebrafish embryos. A) Correlation of Placeholder candidates in sperm and embryos. Pearson correlation R values are displayed and the order was determined by clustering in sup fig 3a. B) Maintenance of Placeholder candidates during sperm-to-embryo transition. Venn diagrams indicating overlap of called peaks comparing sperm (blue), preZGA (green), and postZGA (red) samples. C) Heatmaps for epigenetic features in sperm and embryos. Displayed are dynamic loci where epigenetic marks change during development (top), and promoter regions (bottom). D) Summary of Placeholder maintenance during sperm-to-embryo transition at hypoDNAme regions.

Next, to investigate maintenance of Placeholder, we directly compared the embryonic peak localization for H3K4me1, H2AFV, and H3K14ac to their respective peak locations in sperm. For H3K4me1, H2AFV, and H3K14ac we observed 31%, 84%, and 7% of peak locations were maintained from sperm to preZGA embryos, respectively (Figure 3b). Notably, expansion of H2AFV in preZGA embryos (relative to sperm) occurred either within or proximal to hypoDNAme regions where H2AFV was found in sperm (Figure 3c). As H2AFV locations were largely maintained, and H3K14ac mostly lost, H3K14ac is not a consistent component of Placeholder nucleosomes in embryos, an interpretation reinforced by K-means clustering at dynamic loci (Figure 3c, top panel) and promoter regions (Figure 3c, bottom panel). Thus, a Placeholder nucleosome consisting of H2AFV and H3K4me1 marks DNA hypomethylated loci in both sperm and in preZGA embryos (summarized in Figure 3d).

### Depletion of nuclear H2AFV causes increased DNAme in embryos

To test whether Placeholder antagonizes DNAme in embryos (similar to sperm), we employed loss-of-function approaches by injecting translation-blocking morpholinos targeting *srcap* mRNA into fertilized 1-cell embryos or oocytes, followed by *in vitro* fertilization (IVF). *Srcap* morphants from injected 1-cell embryos died at 3dpf, with phenotypes resembling our genetic *srcap*^−/−^ mutants: an enlarged heart, a recessed jaw, and smaller retinas than WT embryos (Sup Figure 4a-b). Oocyte injections improved morpholino efficacy, partly reducing Srcap protein in preZGA embryos (Sup Figure 4c), which conferred an earlier phenotype involving arrest shortly after ZGA, likely due to maternal mRNA targeting (Sup Figure 4d). Notably, this partial reduction conferred a moderate yet significant increase in DNAme at regions normally marked by H2AFV (Sup Figure 4e), with more significant increases at loci that normally undergo cleavage-phase loss of DNAme (Sup Figure 4e). Thus, Srcap knockdown had a greater effect at regions undergoing both DNA demethylation and H2AFV acquisition. Although these results support H2AFV in deterring DNAme, we sought an orthogonal loss-of-function approach.

The eukaryotic chaperone Anp32e antagonizes H2AFV incorporation by binding and preventing the H2AFV C-terminus from interacting with H3 (Mao et al., 2014; Obri et al., 2014) (Figure 4a). Here, we depleted H2AFV from chromatin via ectopic elevation of Anp32e protein in embryos by injecting purified recombinant Anp32e into 1-cell embryos (Figure 4b). This approach conferred arrest at or prior to early somite stages of development in all injected animals (Figure 4c). Utilizing a transgenic H2AFV:GFP tracer line (Pauls et al., 2001), Anp32e injection caused re-localization of cleavage phase H2AFV from the nucleus to the cytoplasm (Figure 4d). Quantification revealed a 4-fold increase in cytoplasmic H2AFV (Figure 4e), which was confirmed by cell fractionation assays (Figure 4f). Thus, excess Anp32e causes considerable, though incomplete, loss of H2AFV from nuclei. Interestingly, RRBS-Seq on Anp32e-injected embryos revealed a significant increase in DNAme (at 4hpf) at genomic regions which are normally maternally reprogrammed (Figure 4g) - those which are inherited as high DNAme from the maternal allele and become hypoDNAme in embryos - similar to our observation using the Srcap morpholino. Notably, these regions were less CpG rich than non-impacted regions (Figure 4h). Taken together, Anp32e injection caused partial nuclear depletion of H2AFV and increased DNAme largely at weak (CpG poor) and embryonically reprogrammed H2AFV sites.

**Figure 4:**
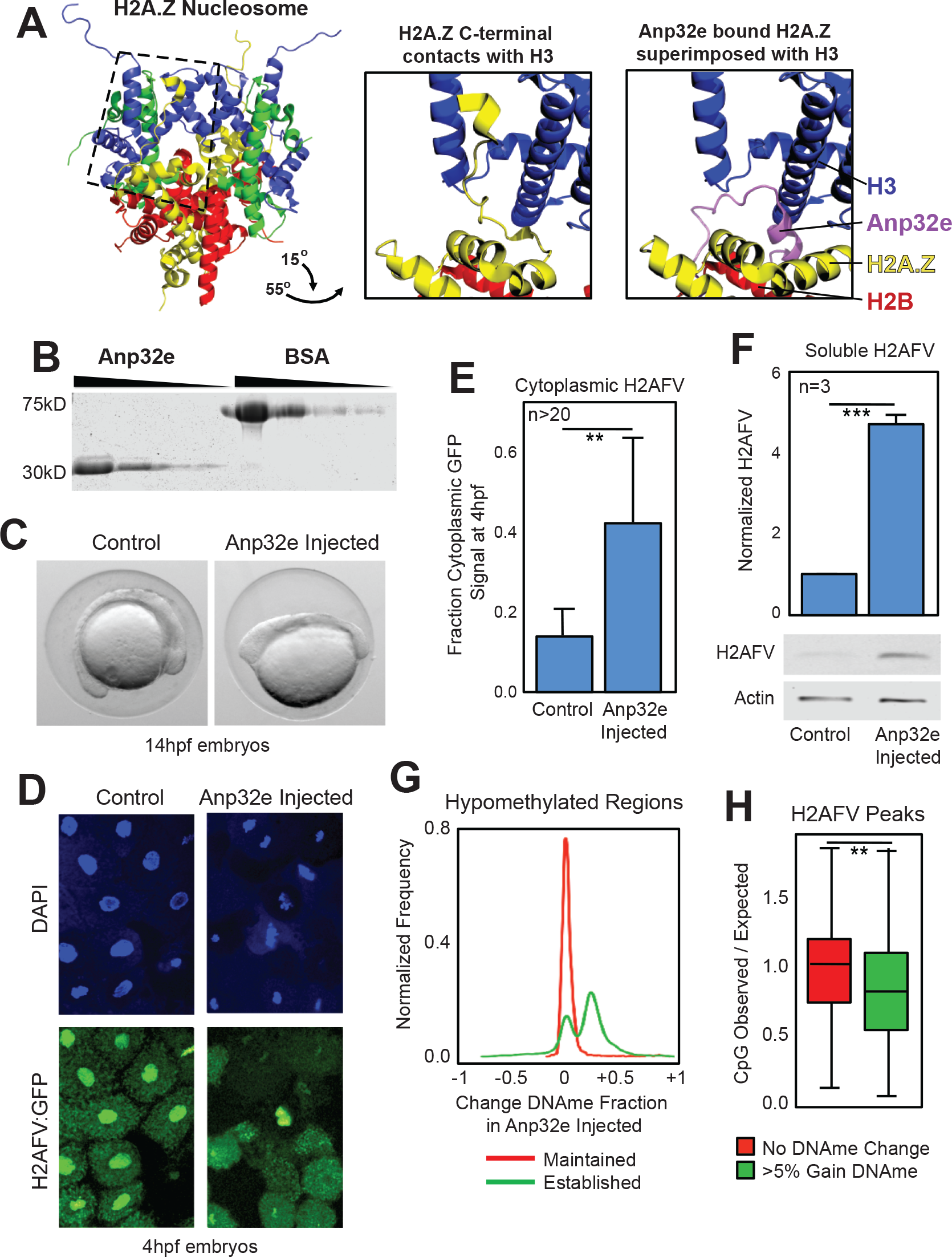
Anp32e protein injection causes reduced nuclear H2AFV and increased DNAme in embryos. A) Anp32e binding inhibits H2A.Z within the nucleosome. Left-crystal structure for the nucleosome octamer containing mouse H2A.Z (yellow). Middle- zoomed in and rotated view of the H2A.Z nucleosome octamer (dashed box from left) where the C-terminus of H2A.Z interacts with histone H3. Right- the same view as middle panel, with the superimposed H2A.Z bound to human ANP32E. The C-terminus of H2A.Z no longer interacts with histone H3. B) Purification of recombinant zebrafish Anp32e protein. A comparison to bovine serum albumin (BSA) demonstrates purity. C) Phenotype of early somite staged zebrafish following Anp32e protein injection. D) H2AFV signal is depleted in nuclei following Anp32e injection in embryos. DAPI and H2AFV:GFP localization in cells from sphere staged (4hpf) embryos. Injection was into the 1-cell embryo. E) Microscopy based quantification of cytoplasmic H2AFV following Anp32e injection. Fraction of H2AFV:GFP signal in control and injected embryos as in panel D. T-tests (see methods): ** = p< 0.01 F) Quantification of H2AFV in cellular fractions resulting from Anp32e injection. Western blotting for soluble H2AFV indicated that Anp32e injection caused increased cytoplasmic H2AFV. T-tests: *** = p < 0.001. G) Anp32e injection causes increased DNAme at reprogrammed loci. Regions were parsed based on the degree of maternal DNA demethylation that occurred in WT embryos - less 0.2 fraction embryonic DNAme and either less than (green) or greater than (red) 0.2 fraction DNAme in oocytes. H) DNAme increases at CpG depleted loci. CpG density (observed/expected) was determined at WT H2AFV sites and parsed based on change in DNAme following Anp32e injection. T-tests: ** = p< 0.01

### Embryos lacking Anp32e have increased H2AFV and decreased DNAme

Next, we conducted the converse experiment to determine whether increased H2AFV in embryos causes DNAme loss. Here, we measured H2AFV and DNAme at sphere stage (4hpf) in WT and three types of *anp32e* mutant embryos: *anp32e* maternal zygotic nulls *(anp32e*^−/−^), *anp32e*^−/+^ maternal null zygotic heterozygotes (Mat-Null), and *anp32e*^+/−^ paternal null zygotic heterozygotes (Pat-Null) (Figure 5a). K-means clustering to compare embryo to sperm enrichment of H2AFV yielded four clusters: Cluster 1 regions are occupied by high H2AFV in both WT sperm and sphere embryos. Cluster 2 regions contain moderate-to-high H2AFV in all samples, with higher levels observed when maternal Anp32e is lacking. Cluster 3 regions lack H2AFV in WT embryos, but contain H2AFV in *anp32e* mutants, and thus appear to rely on Anp32e to antagonize H2AFV. Cluster 4 regions acquire H2AFV in sperm lacking Anp32e, but not in embryos lacking Anp32e. Thus, these loci lose H2AFV in embryos independent of Anp32e. Notably, all three *anp32e* mutants display an increase in H2AFV occupancy, with slightly less H2AFV acquisition in the Pat-Null, as maternally-contributed Anp32e protein or RNA likely provides some H2AFV removal.

**Figure 5:**
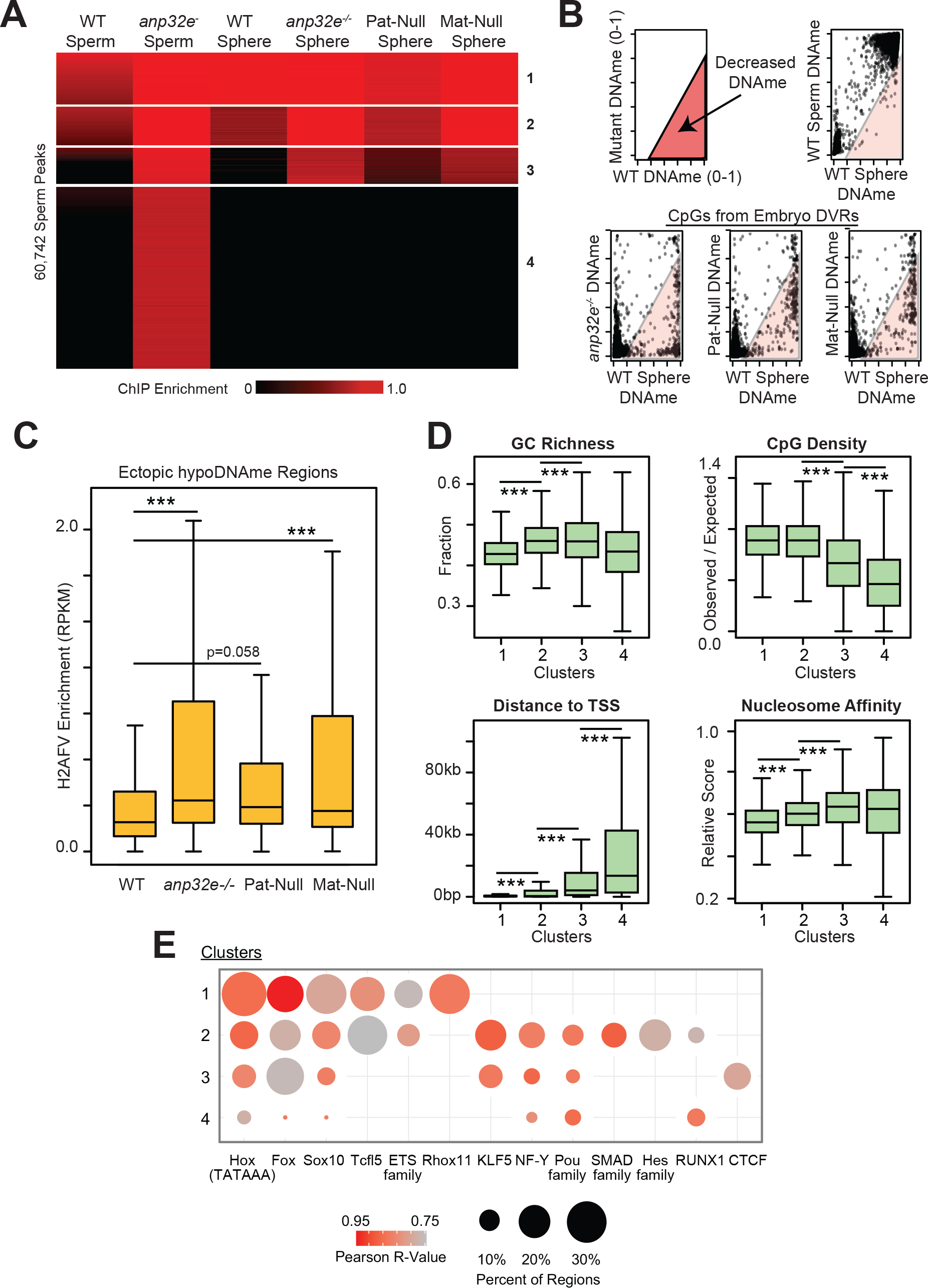
*anp32e* mutation inheritance dictates impact on H2AFV and DNAme. A) ChIP-Seq for H2AFV in sperm and sphere stage embryos comparing WT to *anp32e* mutants. Displayed is a heatmap of K-means clustered Z-scores at all H2AFV sperm peaks. B) Scatterplots of fraction DNAme comparing mutant to WT samples. Top - a diagram for where decreased DNAme can be found (pink shaded area) and an example of maintained DNAme comparing WT sperm to embryos. Bottom - *anp32e* mutant DNAme compared to WT at differential H2AFV regions (DVRs). C) H2AFV increases where DNAme decreases in mutants. Boxplots of H2AFV enrichment are displayed. T-tests (see methods) ***2 = p < 0.001. D) H2AFV increases at CpG depleted high nucleosome affinity sites in mutants. Clusters from panel A were analyzed for GC richness, CpG density, distance to nearest transcription state site (TSS) and predicted nucleosome affinity. T-tests (see methods) *** = p < 0.001. E) Transcription factors motifs differ at ectopic verses maintained H2AFV sites. Clusters from panel A were analyzed using RSAT. Motif frequency is indicated by circle size, and similarity to known binding motifs is indicated by Pearson R values in shaded red.

We then used RRBS to examine whether reductions in DNAme accompanies the increase in H2AFV observed in Clusters 2 and 3. Interestingly, in all three mutant types, DNAme decreased at CpGs within regions where H2AFV increases (Figure 5b, Sup Figure 5a). In keeping, the regions where DNAme was reduced in the mutants gained H2AFV (Figure 5c). Interestingly, the clusters that gained the most H2AFV in embryos (Cluster 3) or sperm (Cluster 4) exhibited lower median CpG density (but similar GC richness) and higher nucleosome affinity (Figure 5d). These results raise the possibility that Anp32e removes promiscuous H2AFV-containing nucleosomes from high-affinity non-promoter regions to help ensure that these regions can acquire proper DNAme (see Discussion).

Next, to identify candidate factors for targeting SRCAP complex, we performed Regulatory Sequence Analysis (Medina-Rivera et al., 2015) and identified enriched TF motifs in each cluster (Figure 5e). Cluster 1 was enriched for Homeobox-and ETS-family TF motifs. Cluster 2 was additionally enriched for Klf5, NF-Y, Pou-family, SMAD-family, Hes-family, and Runx1 motifs. Clusters 3 and 4 showed diminished enrichment for these same factors – along with one unique and strong enrichment, CTCF. Interestingly, H2AFV occupancy was dramatically increased in *anp32e* mutants specifically at differentially methylated CTCF sites (Sup Figure 5b).

### Anp32e loss confers activation at genes that gain H2AFV

As H2A.Z depletion causes differentiation and inappropriate gene expression in mouse cell culture (Hu et al., 2013), we next determined the impact of *anp32e* loss on postZGA embryo gene transcription. Notably, ~7000 genes were misregulated in *anp32e*^−/−^ mutants at 6hpf, compared to ~1600 in Mat-Null or Pat-Null embryos (Figure 6a). Here, Gene Ontology (GO) analysis of upregulated genes yielded the same GO terms with all mutant types (Homeobox, DNA-binding, transcriptional regulation, and developmental proteins), with the statistical significance highest in the homozygous *anp32e*^−/−^ mutant (Figure 6b). (note: downregulated genes were not statistically significant).

**Figure 6:**
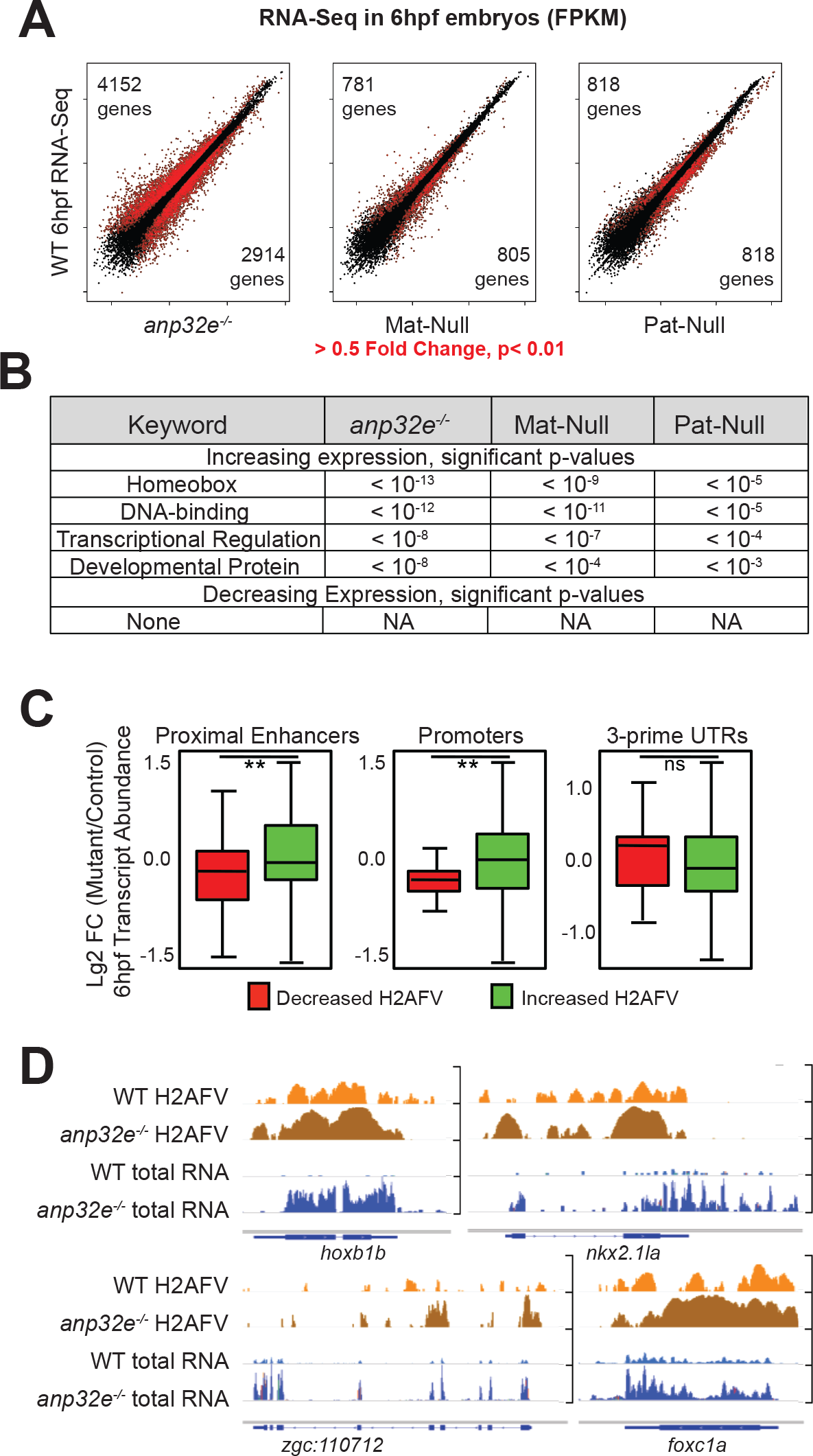
*anp32e* null mutation causes inappropriate early activation of developmental genes. A) RNA-Seq on *anp32e* mutant postZGA embryos (6hpf). Displayed is a comparison of mutant (x-axis) to WT (y-axis) FPKM values, with red indicating expression changes greater than 0.5-fold (either increasing or decreasing) in mutant verses WT samples. The number of genes with increasing expression is indicated in the top left corner and decreasing gene number is in the bottom right corner. B) Developmental genes are mis-regulated in *anp32e* mutants. DAVID Gene Ontolgoy characterized biological function. C) Increased H2AFV at promoter and enhancer regions correlate with increased proximal Hox-gene transcription. Strategy for parsing changes in H2AFV is displayed in supplemental figure 6. T-tests (see methods): ** = p< 0.01, ns = not significant. D) A gallery of browser snapshots to illustrate increased H2AFV is associated with increased gene expression at *hox* family genes. ChIP-Seq data is Log10FE (0-1) values, and RNA-Seq is normalized FPKM.

Thus, excess H2AFV correlated with increased expression at developmental genes, especially *hox* genes. As expected based on the *anp32e*^−/−^ adult phenotype, these expression differences resolve as development progresses (Sup Figure 6a). To investigate whether this upregulation was a direct result of H2AFV accumulation, we compared our H2AFV ChIP profiles in *anp32e*^−/−^ embryos with changes in proximal gene expression (thresholding in Sup Figure 6b). Notably, increased H2AFV at promoters and enhancers (but not 3’-UTRs) correlated with increased expression (Figure 6d). A set of browser screenshots at several affected genes illustrates this (Figure 6e). Taken together, loss of Anp32e increases H2AFV at the promoters and enhancers of particular developmental regulators (e.g. *hox* genes), which leads to their precocious expression following zygotic genome activation.

## Discussion

A key issue in germline-to-embryo epigenetics in vertebrates and mammals is whether and how parental DNAme patterns are maintained or reprogrammed after fertilization, through the transcriptionally-quiescent portions of cleavage phase. The answers and mechanisms that address this have important implications for developmental potential (totipotency or pluripotency) and transgenerational inheritance. Zebrafish and mammals lack DNAme at particular gene classes in their paternal germlines, including germline-specific genes (e.g. meiosis-specific), housekeeping genes – and interestingly, also at hundreds of TFs important for embryo development, but not germline development (Hajkova et al., 2008; Hammoud et al., 2009; Jiang et al., 2013; Potok et al., 2013; Seisenberger et al., 2012; Vincent et al., 2013). These gene classes are packaged in two distinctive types of chromatin in zebrafish sperm: the ‘active’ meiosis and housekeeping gene promoters are packaged in chromatin containing H3K4me1/2/3 and acetylation, whereas the ‘silent’ developmental promoters are packaged in bivalent chromatin (H3K4me3 with H3K27me3) with H2AFV, and lack acetylation. However, these marks are not simply fully retained from the gametes into the early embryo (Akkers et al., 2009; Dahl et al., 2016; Liu et al., 2016; Vastenhouw et al., 2010). Therefore, a key issue is how the DNA methylation patterns of the sperm are maintained in the embryo during a period that lacks many of the known deterrents of DNAme. Likewise, regarding the maternal genome, zebrafish oocytes also lack DNAme at housekeeping genes and a portion of developmental genes (co-incident with high H3K4me3) – but in contrast to sperm, oocytes bear DNAme at hundreds of loci important for embryo development and germline identity. Whereas the DNAme patterns in zebrafish sperm are virtually identical to pluripotent sphere stage embryo patterns, the maternal genome is reprogrammed gradually during cleavage such that DNAme patterns reflect the paternal genome (Jiang et al., 2013; Potok et al., 2013). Here, our combined genomic and functional evidence demonstrate that a Placeholder nucleosome underlies these observations – to harmonize the DNAme patterns of the two parental genomes, and to provide proper transcriptional regulation in early embryos.

We begin by discussing the conceptual issue – why utilize a Placeholder nucleosome (bearing H2AFV and H3K4me1) during the germline-to-embryo transition to passively deter DNAme – why not simply retain the parental marks? First, since the epigenetic/variant marking of the two parental genomes is highly different, retaining parental marks does not allow for harmonization of the two parental epigenomes in the embryo. Second, as zebrafish do not express TET proteins (or AID) during cleavage stage, no known mechanism for targeted active demethylation appears present. Moreover, since cleavage lasts ten cell/replication cycles, a nucleosome that passively deters DNAme presents a simple and logical alternative. Finally, although H3K4me3 (or bivalency) deters DNAme, H3K4me3 is typically linked to transcription initiation; as cleavage stage generally lacks transcription, transcription-independent surrogate marks are needed to deter DNAme. Thus, the Placeholder nucleosome provides a mechanism both for maintaining the vast majority of the paternal regions lacking DNAme, while also enabling reprogramming of maternal DNAme patterns. Regarding the name ‘Placeholder’ – during cleavage phase Placeholder occupies DNA hypomethylated genes ‘in-place-of’ the other regulatory chromatin, which at ZGA will resolve into either active (housekeeping genes) or poised/silent (developmental genes) chromatin regions, both of which are DNA hypomethylated.

Although the reprogramming of paternal DNAme patterns is different between zebrafish and mammals (which reprogram via TET proteins), the maternal mammalian genome is demethylated in a similar manner to zebrafish – by passive mechanisms spanning ~8 cell cycles (Amouroux et al., 2016; Smith et al., 2012). One intriguing possibility is that mammalian maternal DNA demethylation also relies on Placeholder-type nucleosomes. Here, it will be of high interest to determine whether a Placeholder type of nucleosome (with H2A.Z and/or H3K4me1, or alternative marks that deter DNAme) also marks and regulates the passive DNAme reprogramming on the maternal genome during early mammalian development. Furthermore, as H3K4me manipulation affects trans-generational inheritance (Siklenka et al., 2015), H3K4me alterations on discrete Placeholder nucleosomes may facilitate this passage. Finally, interplay may exist between the two marks that define Placeholder: H2AFV and H3K4me1. H3K4me1 can inhibit DNMT function *in vitro* (Guo et al., 2015), and H2A.Z promotes the recruitment of the MLL complexes – the dominant family of H3K4 methyltransferases (Hu et al., 2013). Importantly, mRNAs encoding orthologs of all four MLL-type complexes (1-4) are present in zebrafish embryos (data from Lee et al., 2013), raising the possibility of a positive feedback loop between H2AFV and H3K4me1 that collaborates to restrict DNMT function in embryos.

Our functional studies involved modulating Srcap, the H2AFV installation remodeler, and Anp32e, the H2AFV-specific removal chaperone. Here, either Srcap depletion or Anp32e elevation conferred H2AFV loss and increased DNAme in embryos. Conversely, Anp32e loss resulted in H2AFV acquisition (largely within H3K4me1-marked intergenic regions) and decreased DNAme in sperm and early embryos – which together provide the functional evidence that H2AFV, often in combination with H3K4me1, deters DNAme. Notably, H2AFV gain in *anp32e* mutants occurs at loci/sequences with high nucleosome affinity and low-intermediate CpG content. We suggest that H2AFV nucleosomes are more stable at high-affinity sites than their canonical counterparts – and that the normal function of Anp32e is to ‘prune’ H2AFV from the genome to avoid accumulation at high-affinity sites that lack constant Srcap installation (Placeholder antagonism, Figure 7). This pruning may ensure high H2AFV (H2A.Z) occupancy only at sites engaging in consistent active incorporation by Srcap complex (Placeholder establishment, Figure 7), and not at adjacent low-affinity sites – which may explain the striking focal co-incidence of H2AFV and hypoDNAme. Indeed, accumulation of H2AFV might be predicted to cause precocious expression of genes, especially those of low-intermediate CpG content, perhaps by lowering DNAme and enabling TF binding. In keeping, we observed a coordinate premature increase in transcription of particular genes that accumulate H2AFV in *anp32e* mutants, such as *hox* genes. Thus, Anp32e refines H2AFV occupancy to ensure that precocious transcription is minimized and that ZGA occurs in the most precise manner.

**Figure 7:**
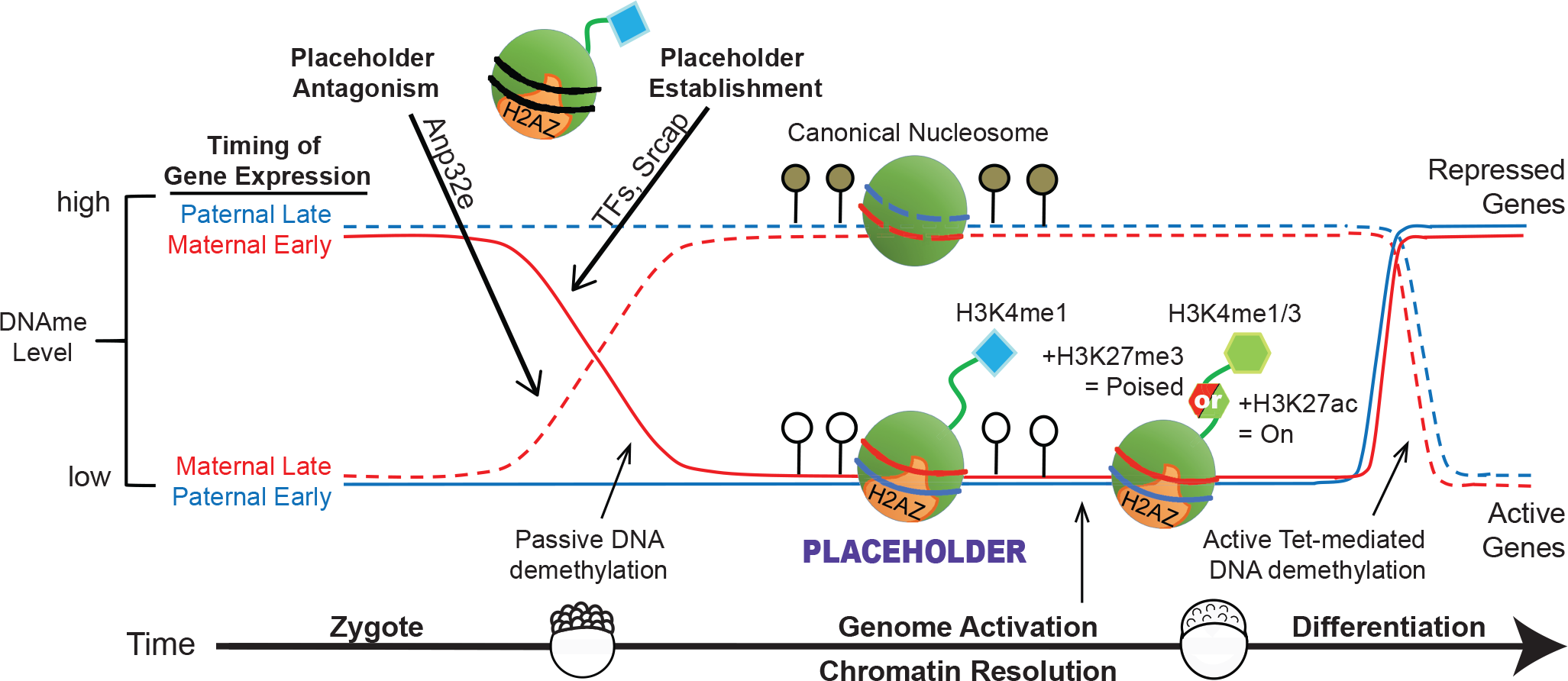
Model for Placeholder regulation of DNAme reprogramming. DNAme reprogramming during cleavage phase relies on both establishment and antagonism of Placeholder nucleosomes containing H2AFV and H3K4me1 to enable subsequent transcriptional activation. Placeholder is established at maternal regions which gradually undergo demethylation, and Placeholder is antagonized at maternal regions where default acquisition of DNAme occurs. Concurrent with zygotic transcriptional activation, bivalent marks are established.

Finally, our identification of TF motifs enriched at Placeholder-occupied sites provides candidate TFs for Srcap complex recruitment, and possibly also MLL3/4 orthologs (which catalyze H3K4me1), to establish Placeholder attributes (Figure 7). Among the identified motifs was CTCF, where H2A.Z accumulated specifically in *anp32e* mutants, consistent with a prior study in mammalian cells (Obri et al., 2014). Thus, dynamic recruitment/removal of H2A.Z/FV may occur at CTCF motifs to impact the DNAme status and CTCF binding – potentially altering chromatin looping. More broadly, candidate Srcap recruiting factors during cleavage stage include proteins in particular families (Hox, Ets, Klf, Pou, Fox, Smad, Hes) as well as NF-Y and Runx1. Notably, genes fated for a ‘poised’ state (bivalency) are more enriched for Hox-, Fox- and Pou-family factors, whereas those for housekeeping genes were more enriched for Ets-family factors – suggesting specialization for gene classes. Here, we speculate a sequential ‘hand-off’ of cooperative TFs: first, TFs impose H3K4me3 or bivalency in the germline to deter DNAme at particular loci in gametes. Second, during cleavage phase (transcriptional quiescence), a largely separate set of TFs (described above) recruit Srcap complex to install Placeholder (Placeholder establishment, Figure 7) – providing a mechanism to define all DNA hypomethylated loci, which ‘funnel’ through a common chromatin Placeholder chromatin state prior to ZGA. Lastly, during and after ZGA, cell-type specific TFs resolve Placeholder loci into either active or bivalent loci status, through their selective recruitment of additional chromatin modifying factors. Taken together, our work provides evidence for the concept that DNAme status can be inherited from the gamete and maintained in the embryo through the installation of a surrogate chromatin Placeholder during transcriptional quiescence.

### Genomics and Data Access

Data may be downloaded from GEO with accession GSE95033

**Table 1.** Antibodies Used

| Target | Company - Catalog number | Use |
| --- | --- | --- |
| H2A.Z | Active Motif - 39113 | ChIP |
| H3K4me3 | Active Motif - 39159 | ChIP |
| H3K4me1 | Active Motif - 39297 | ChIP |
| H3K27ac | Abcam - ab4729 | ChIP |
| H3K27me3 | Active Motif - 39155 | ChIP |
| H3K14ac | Active Motif - 39599 | ChIP |
| Srcap | Abcam - ab99408 | Western |
| Histone H3 | Abcam - ab46765 | Western |
| H2A.Z | Abcam - ab4174 | ChIP |
| H3K4me1 | Abcam- ab8895 | ChIP |
| Actin | Custom from Shaw Lab (Utah) | Western |

## Acknowledgements

We thank Brian Dalley for sequencing expertise, and Tim Parnell, Tim Mosbruger, and Jingtao Guo for discussions regarding bioinformatics. We thank Jason Gertz, Margaret Kasten, Cedric Clapier, Simon Curie, and Edward Grow for advice on techniques. We extend a special thanks to KT Varley for helping us to adapt reduced representation bisulfite sequencing, to David Grunwald and his laboratory for teaching us germline transplantation, and to Marnie Halpern for generously sharing the H2AFV:GFP fish. B.R.C. is an investigator with the HHMI. Financial support was from Howard Hughes Medical Institute (HHMI) (genomics, biologicals), and CA24014 for Huntsman Cancer Institute core facilities. P.J.M. was funded by the Eunice Kennedy Shriver National Institute Of Child Health & Human Development of the National Institutes of Health under Award Number T32HD007491. Imaging was done at the University of Utah Fluorescence Microscopy Core Facility grant 1S10RR024761-01.

### Author Contributions

Experimental design: B.R.C., P.J.M., and S.F.W; experiment execution: P.J.M., S.F.W., C.R.J, and C.L.W.; data analysis and figures: P.J.M; manuscript preparation: P.J.M and B.R.C with comments from all authors.

---

**Supplemental Figure l.**
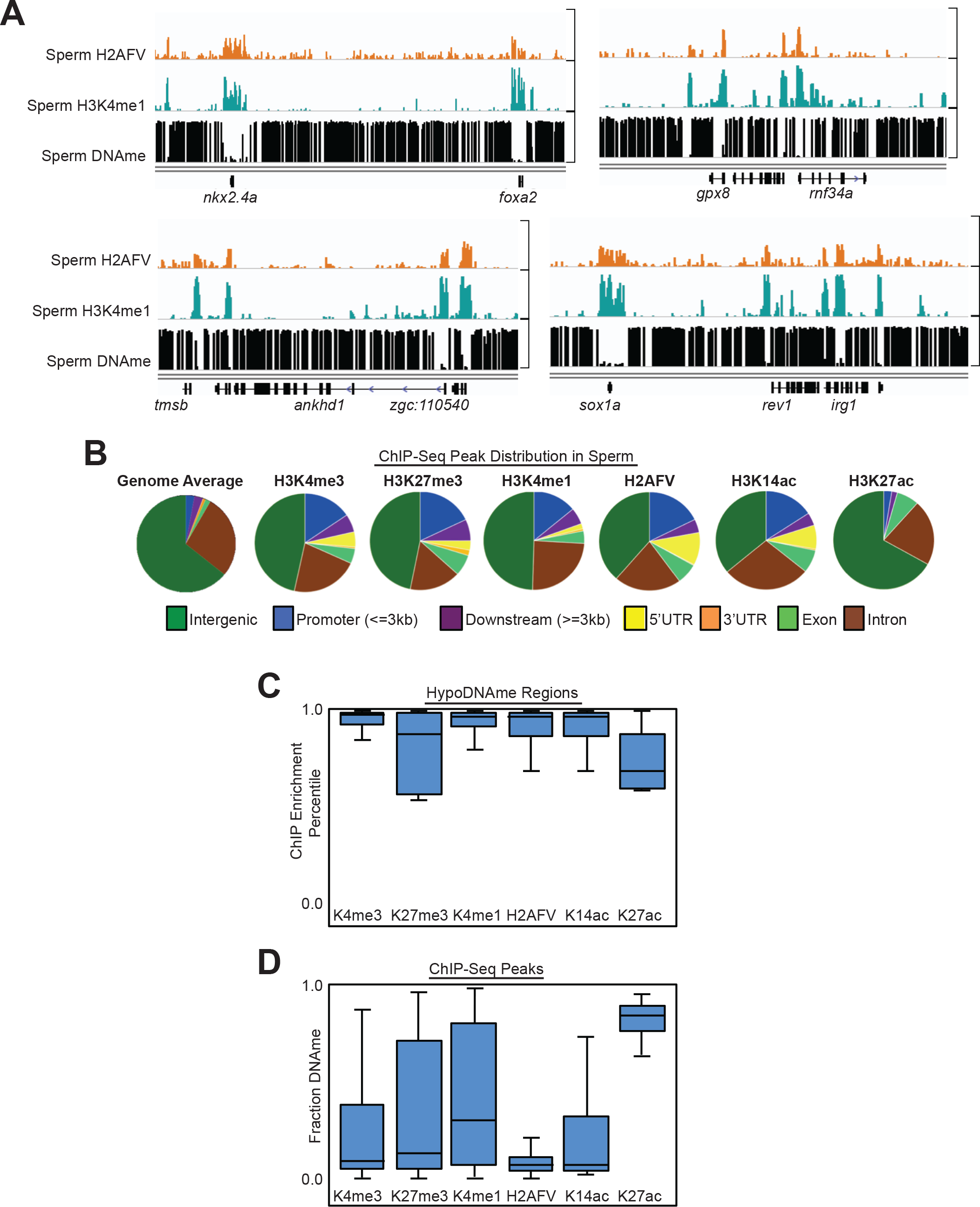
A) A gallery of browser snapshots displaying enrichment for H2AFV, H3K4me1, and DNAme in sperm. DNAme data from Potok et al. 2013. Log10FE (0-1) values for ChIP-Seq data and DNAme fraction (0-1). B) Distribution of ChIP-Seq peaks in sperm assaying for various chromatin marks across a series of genomic landmarks. Non-enriched genome wide distribution is displayed for comparison. C) ChIP enrichment percentiles for various chromatin marks at hypoDNAme regions in sperm. D) DNA methylation level (fraction) for enriched regions assays by ChIP-Seq in sperm.

**Supplemental Figure 2.**
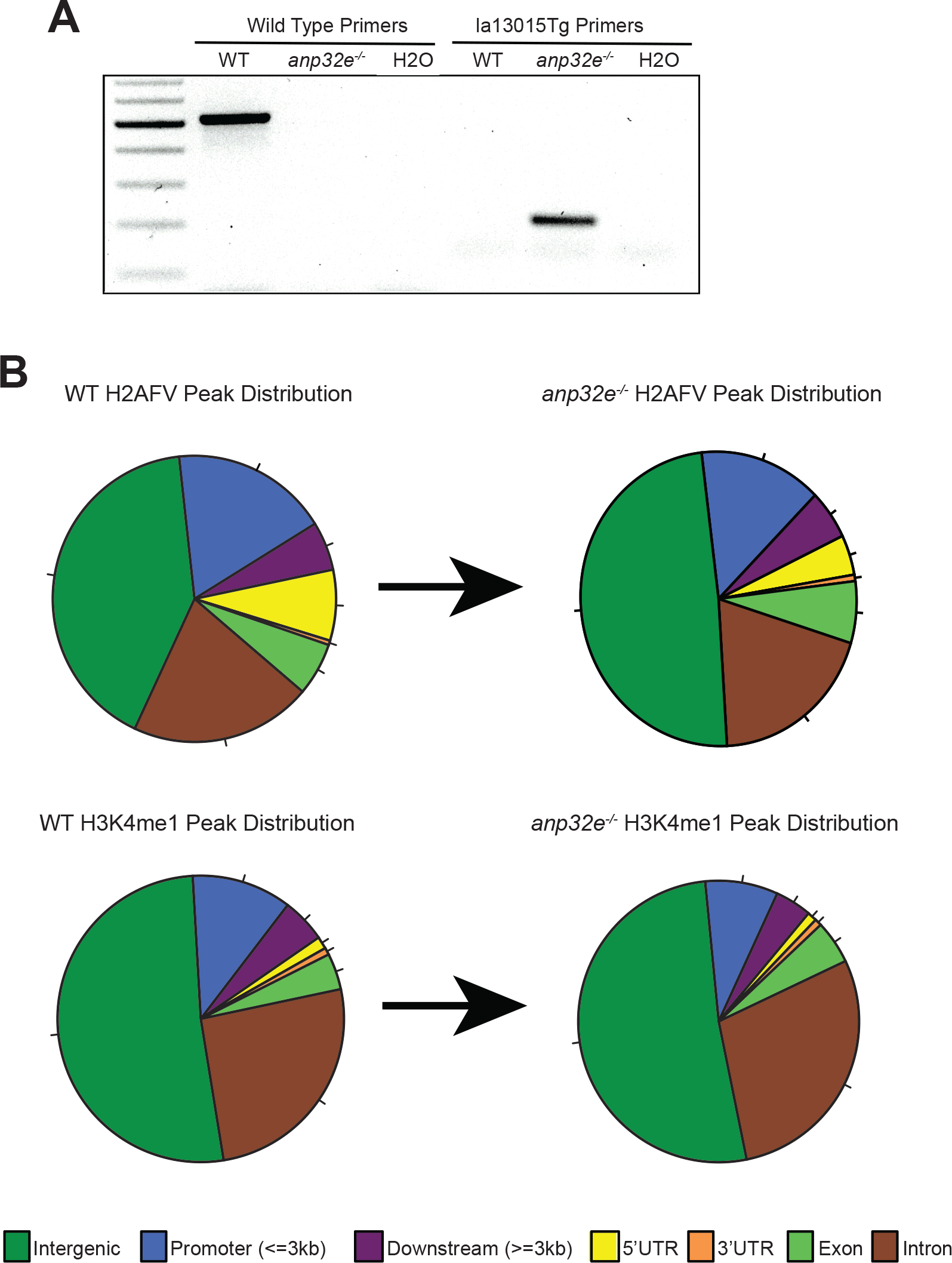
A) Electrophoresis gel of PCR products from genotyping of la13015Tg insertion. Homozygous *anp32e* mutant fish have only the smaller mutant band and not the wild type band. B) Change in peak distribution (as in supplemental figure 1a) for H2AFV and H3K4me1 in *anp32e* null sperm compared to wild type.

**Supplemental Figure 3.**
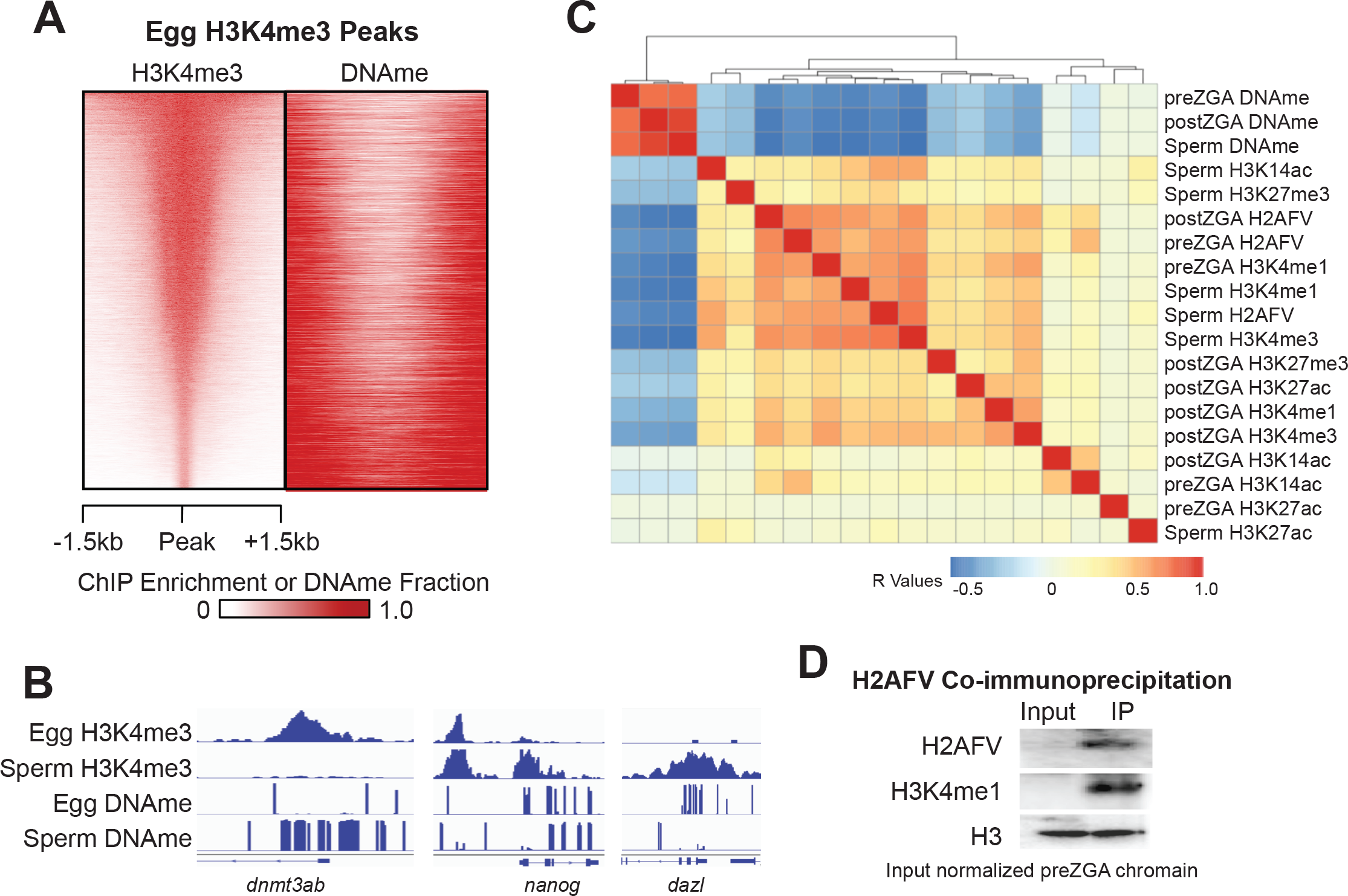
A) Heatmaps of H3K4me3 and DNAme at H3K4me3 peaks in eggs indicating the two epigenetic marks are anti-correlated with one another. B) Browser snapshots of enrichment for H3K4me3 and DNAme in sperm and eggs. H3K4me3 marks many promoters, but is absent from the promoters of transcription factor genes in eggs which also have moderate to high DNAme in eggs. Normalized read depth for ChIP-Seq data and DNAme fraction (0-1). C) Clustering of pairwise Pearson correlation R values for all epigenetic features assayed in sperm, and in pre- and post-ZGA embryo samples. H2AFV and H3K4me1 are most strongly correlated with one another and anti-correlated with DNAme. D) H3K4me1 co-immunoprecipitates with H2AFV in preZGA embryos. Lysates were chromatin from preZGA embryos, and western blot loading was normalized to input histone H3 level.

**Supplemental Figure 4.**
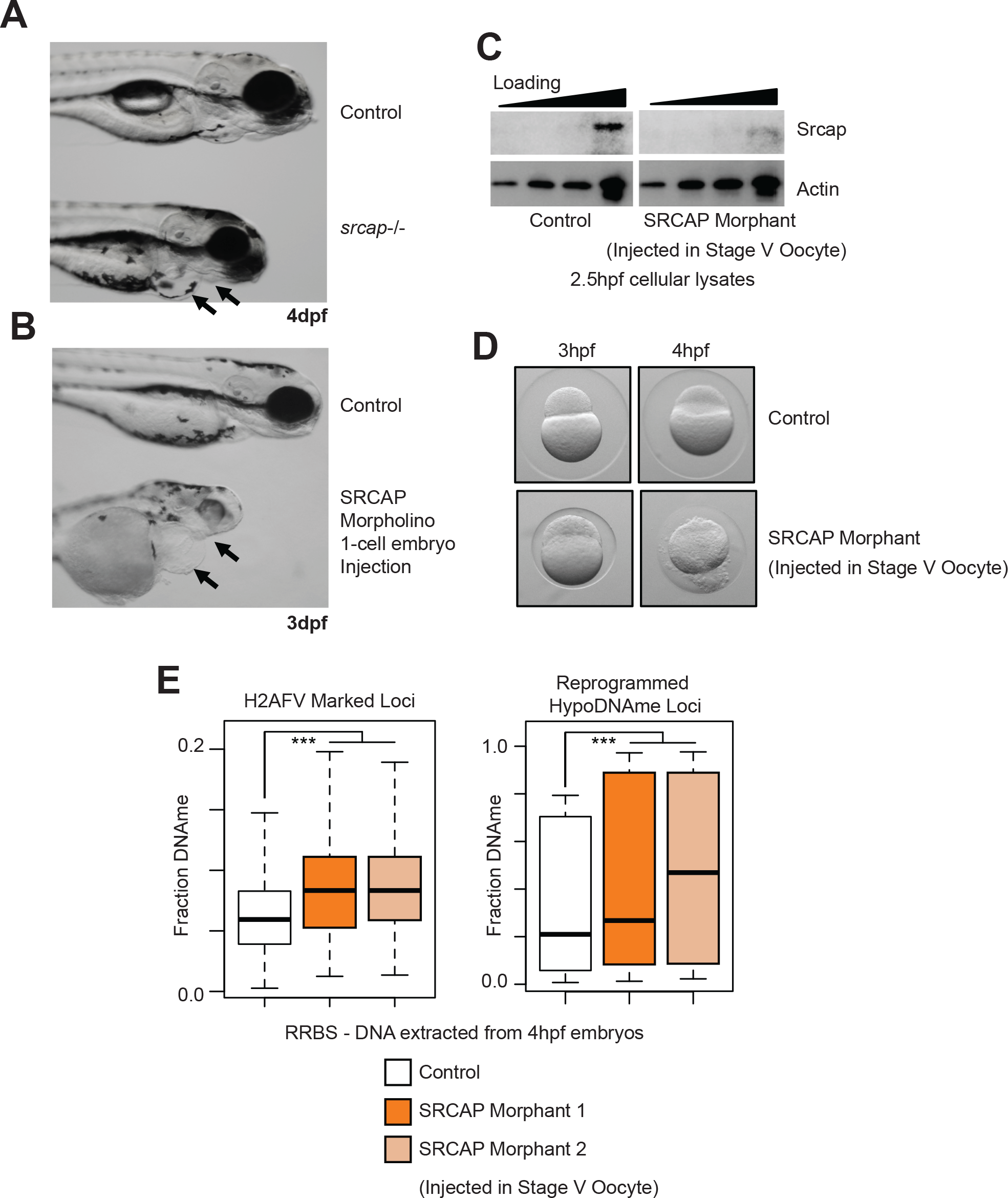
A) Phenotype of zygotic null *srcap*^−/−^ mutant fish compared to wild type at 4 days post fertilization. Arrows indicate recessed jaw, and enlarged heart in mutant embryos. B) Similar to panel A, except phenotypic characterization of embryos injected with an SRCAP morpholino. Embryos are at 3dpf. As in panel A, arrows indicate a recessed jaw and enlarged heart. C) A western blot for Srcap abundance comparing cellular lysates from control 2.5hpf embryos to lysates from embryos that resulted from *in vitro* fertilization of oocytes injected with an *srcap* morpholino. D) *srcap* morphants from stage V oocyte injection are modestly impaired at 3hpf, but have gross morphological defects at 4hpf – immediately following zygotic genome activation. E) RRBS-Seq DNAme analysis of *srcap* morphants compared to control embryos for all wild type H2AFV marked loci (left), and for loci which undergo DNA demethylation on the maternal allele under wild type conditions.

**Supplemental Figure 5.**
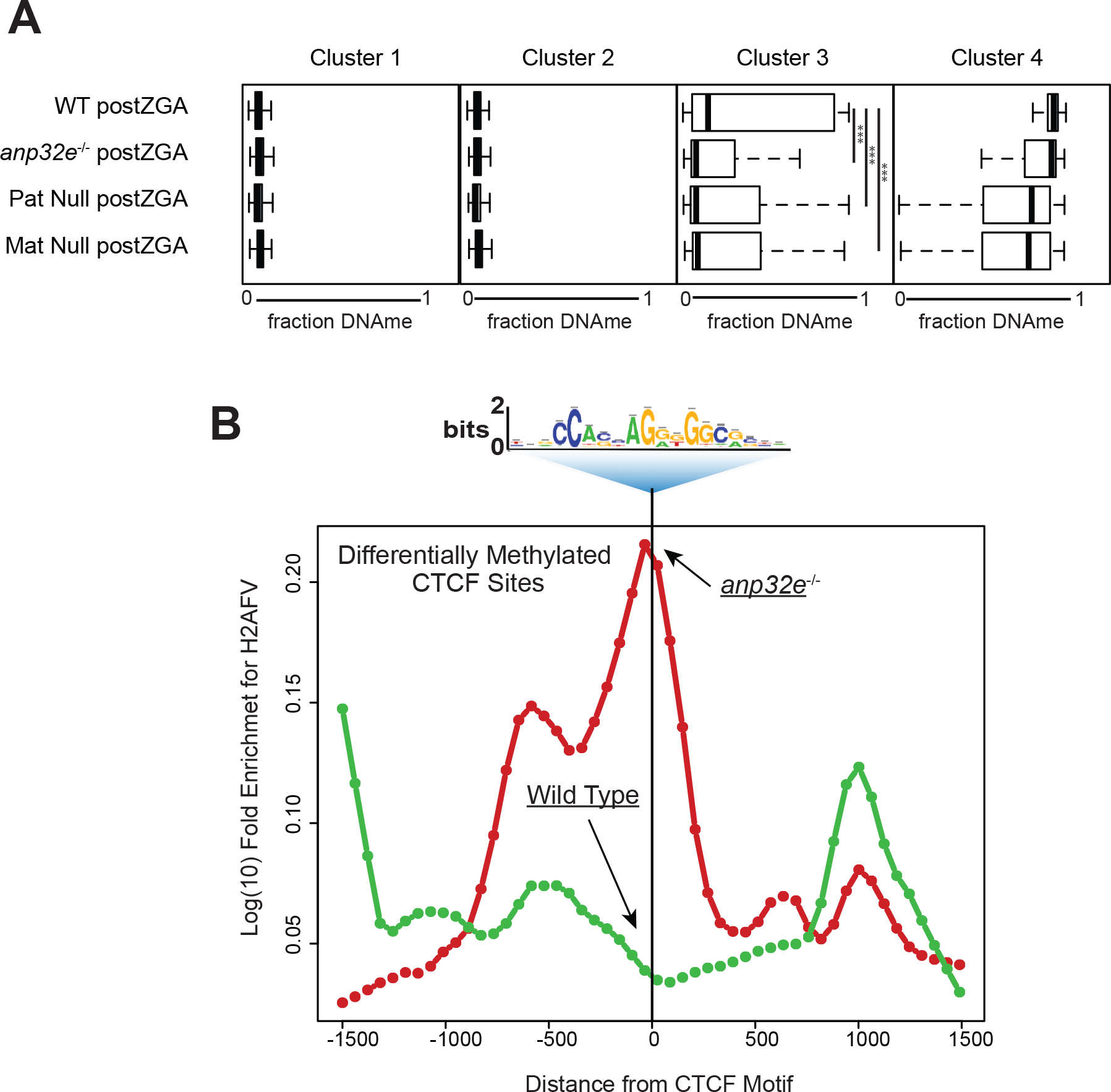
A) Boxplots for DNAme in embryos (labeled on the left) parsed based on K-means clustering from figure 5a. x-axis is fraction DNAme and *** = p < 0.001. B) H2AFV is significantly higher in the *anp32e*^−/−^ embryos than in the wild type control embryos at differentially methylated CTCF sites (p<0.001). CTCF motifs were identified using RSAT analysis applied to regions from figure 5a cluster 3. These motifs were then mapped to the genome and CTCF containing regions were parsed based on DNAme change in the *anp32e* null embryos.

**Supplemental Figure 6.**
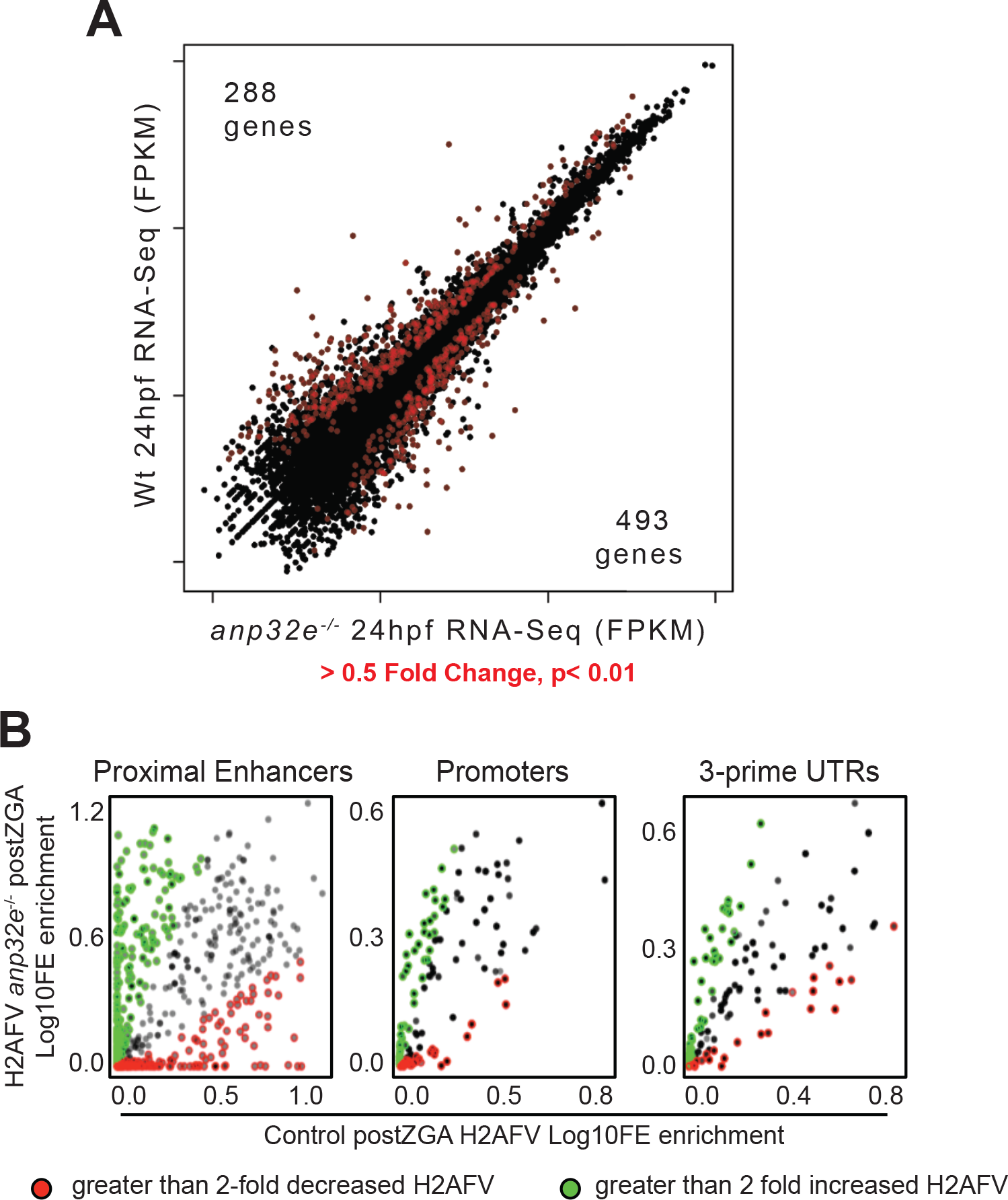
A) RNA-Seq was performed on *anp32e* mutant embryos (24hpf) and compared to wild type. Values for every gene in the genome with mutant data is on the x-axis and wild type on the y-axis (FPKM), and red indicating expression changes greater than 0.5-fold (either increasing or decreasing). The number of genes with increasing expression is indicated in the top left corner and decreasing gene number is in the bottom right corner. B) Log10 Fold Enrichment for H2AFV was measured for previously described ChIP-Seq data (Figure 5) in *anp32e* mutants at three types of genomic features within *hox* gene regions (top GO-term). Feature types were proximal enhancers (marked by H3K27ac and H3K4me1 and within 5kb of a gene), promoters, or 3’-UTRs. These regions were then classified based on whether there was greater than a 2fold change in H2AFV enrichment. Increasing H2AFV regions are marked in green and decreasing in red.

## Methods

### ChIP-Seq

Sperm ChIP-Seq methods are as described previously (Hammoud et al. 2009 and Wu SF et al. 2011). For ChIP in early embryos, whole embryos were cross-linked with 2.2% formaldehyde for 15 minutes before quenching with 250mM glycine. Nuclei were then purified as described previously (Potok et al. 2013) and re-suspended in IP dilution buffer (16.7mM Tris-HCL, 167mM NaCl, 1.2mM EDTA, 1.1% Triton-X, 0.1%SDS). Lysates were then sonicated using a Branson Digital sonicator at 30% amplitude for 8 cycles of 10-pulse sonication (1 pulse per second) with 30sec rest on ice between each round of 10-pulse sonication. 5ug of antibody was then added to the lysate and left overnight to bind chromatin at 4C rotating. The following morning Protein A/G Dynabeads (ThermoFisher ID: 10003D) were bound to the lysate at 4C for 4 hours. Samples were then washed 4 times in RIPA buffer (50mM HEPES, 500mM LiCl, 1mM EDTA, 1% NP-40, 0.7% SDS) and eluted overnight at 65C in RIPA. DNA was then purified from eluate by phenol-extraction and alcohol precipitation. Libraries were made using the NEBNext ChIP-Seq Master Mix Set (New England Biolabs ID: E6240S). High-throughput sequencing was by Illumina’s protocol for 50bp single-end runs on an Illumina HiSeq 2500. See supplemental table for list of antibodies used.

### RRBS-Seq

100-500ng of DNA was purified from zebrafish embryos and digested with MspI then end repaired using Klenow -exo (New England Biolabs ID: R0106S and M0212S). Methylated barcodes (BioO Scientific ID: 511912) were then ligated to each sample using T4 DNA ligase (New England Biolabs ID: M0202S), then bisulfite treated according to manufacturer’s protocol (Zymo Research ID: D5005). Following bisulfite treatment libraries were PCR amplified for 20cycles using Platimun Taq (ThermoFisher ID: 10966018) and the standard primers and protocol from BioO Scientific methylated barcodes kit. High-throughput sequencing was by Illumina’s protocol for 50bp single-end runs on an Illumina HiSeq 2500.

### RNA-Seq

RNA was purified by trizol-chlorophorm extraction (ThermoFisher ID: 15596026) of whole embryos followed by column cleanup. Briefly, 20-200 embryos (for 4dpf and 6hpf respectively) were lysed in 500ul of Trizol, and following chloroform extraction, the RNA-containing aqueous phase was mixed 1:1 with 100% ethanol and bound to an RNAeasy column (Qiagen ID: 74104) by centrifugation according to manufacturer’s recommendations. Column bound RNA was then washed with RW1 and RPE buffers and then eluted with water. Purified RNA was then prepared as libraries for sequencing using the TruSeq Stranded Total RNA Library Prep Kit with Ribo-Zero (Illumina ID: RS-122-2201). High-throughput sequencing was by Illumina’s protocol for 50bp single-end runs on an Illumina HiSeq 2500.

### Bioinformatics Analysis

Fasta files were aligned to Zv10 using the most updated version of Novocraft Novoalign with standard parameters for ChIP-Seq and RNA-Seq, and using bisulfite mode for the RRBS-Seq experiments. Aligned ChIP-Seq data was then processed using MACS2 for enrichment scoring and peak calling. Briefly, read count normalization was performed on alignment files (BAM) to account for sequencing depth differences then background corrected based on input. Finally, Log10-fold enrichment over input lambda was then calculated for each nucleotide across the entire genome. These scores were then used for peak calling and for downstream bioinformatics analysis, which was performed in R using standard methods – only the ‘pheatmap’ package was used to generating images of clustered Pearson correlation heatmaps. All other clustering was performed using CLUSTER 3.0 and heatmaps were generated using Java TREEVIEW. Regions were classified as peaks based on greater than 3fold enrichment of input lambda, and Differential Variant Regions (DVRs) were defined as regions where HA.Z enrichment increased greater than 2fold in Anp32e sperm compared to wild type. Metagene analysis and regional enrichment profiling were performed using DANPOS. Relative peak enrichment was performed using CEAS. Identification of transcription factor motifs was performed with RSAT in ChIP-Seq mode. For RNA-Seq alignment files, splice junction were converted to genomic coordinates and low quality and non-unique reads were further parsed using Sam Transcriptome Parser (USeq; v8.8.8). Stranded differential expression analysis was calculated with Defined Region Differential Seq which utilizes DESeq2 and the reference Zv10 Ensembl gene table downloaded from UCSC. Aligned RRBS-Seq data was processed using the USeq pipeline for bisulfite converted DNA as described previously (Potok et al. 2013), and only regions with greater 8 reads and 3 CpGs were considered.

### Isolation of pellet and supernatant fractions

The method was adapted from (Wysocka J et al, 2001) with the differences detailed below. Approximately, 100 embryos were dechorionated with pronase (1mg/ml, Sigma, cat# 11459643001), resuspended in deyolking buffer (55mM NaCL, 1.8mM KCl, 1.25mM Sodium Biocarbonate) followed by centrifugation at 500 × g, 4°C for 3mins to collect cellular portion of the embryo. Extraction of detergent soluble proteins from cell pellet was performed by treatment with 200μl Buffer A (10mM Hepes, pH 7.9, 10mM KCl, 1.5mM MgCl2 0.34M Sucrose, 10% Glycerol, 1mM DTT, 0.1% Triton X-100, and protease inhibitor). Cells were incubated on ice for 8mins, then100μl was removed for Total sample. The soluble and pellet fractions were collected by centrifugation at 1300 × g, 4°C for 5mins. 4xSB (SDS containing sample loading buffer) was added to the soluble fractions and boiled at 100°C for 5mins. The pellet fraction was resupsended in equal volume of 100μl of Buffer B (3mM EDTA, 0.2mM EGTA, 1mM DTT, protease inhibitor). To solubilize the chromatin, both pellet and total fractions were incubated with 1000U of Micrococcal Nuclease (NEB, Cat# M0247S) at 37°C for 30mins and finally 4×SB (SDS containing sample loading buffer) was added and the samples were boiled at 100°C for 5mins. Samples were further analyzed by western blot analysis.

### Live Cell/Embryo imaging

Approximately 5 H2AZ:GFP embryos per batch were dechorionated with pronase (1mg/ml, Sigma, cat# 11459643001), for 5mins. The dechorionated embryos were rinsed with a 1:1 mix of 1×PBS and 1mg/ml DAPI and placed on a glass bottom microscope slide dish with approximately 10uL of 1:1 mix 1xPBS and 1mg/ml DAPI. Cells were imaged on Nikon A1 Ti-E inverted microscope equipped with Four Photo Multipliers Tube (PMT) detector unit. Images were taken utilizing 405nM diode laser (DAPI) and 488nm Argon Gas laser (GFP) utilizing a 60x oil immersion objective. Z-sections were acquired for a plane of cells within an embryo at 0.5 μm steps. Intensity measurements were calculated using Image J.

### Mutant generation and genotyping

Germline transplantation assays occurred as described previously (Ciruna et al., 2002). Genotyping of *anp32e* mutant fish acquired from Zebrafish International Resource Center occurred by PCR amplification using two separate PCR reactions, for the wild type allele primers were CGCAGCAATAACCAACCAAATGGA and TCCTGCTGGAAGCGGGAACAATATC, and for the mutant allele primers were CGTCAGCGGGGGTCTTTCAT and TCCTGCTGGAAGCGGGAACAATATC. Mutation in the *srcap* gene were achieved by TALEN and genotyping of the mutant allele (which occurred within exon 13) was by PCR HRMA assays as described previously (Dahlem et al., 2012) using the following primer set: CTGGACCAAACCCAATGCTTT, GGTACCTCC AAGACTTGCGT.

### Anp32e protein purification

The mRNA sequence for zebrafish Anp32e with a C-terminal 10X histidine tag was cloned into the pET28a plasmid, which was then electroporated into BL21-CodonPlus bacteria. Bacteria were then grown at 37C in 3L of LB media until 0D600 reached 0.8. Then, cultures were induced with 0.2mM IPTG and left at 16C shaking overnight to allow for protein expression. The following morning bacteria was gently pelleted, solubilized in lysis buffer (50mM Tris-HCL, 500mM NaCl, 10% glycerol, 0.1% triton-X, 1.5mM BME), and then sonicated using a Misonix sonicator at 50% power for 5 cycles of 1 min on/off. Samples were then pelleted at 20,000g for 10min, and the soluble portion was recover and bound to Ni-NTA resin at 4C for 2 hours. Ni-bound protein was then washed 6 times using 1mL lysis buffer supplemented with gradually increasing immidazol concentrations (5-60mM), then eluted with 4-1mL fractions of lysis buffer containing 200mM immidazol. Protein aliquots of 0.5mg/mL were stored at -80C in 25% glycerol, and diluted 1:5 into water immediately before use.

### Oocyte or embryo injection and inhibitor treatment

Embryo injections for Srcap morpholinos (GATCCTTCCGCCATAGCGGTTTGCC) were as described previously (Rai K. et al., 2006), and Anp32e protein injection occurred similarly except 0.1ng of protein was injected into the yolk of each 1-cell embryo. Oocyte injection and *in vitro* fertilization were performed using Srcap morpholino concentrations identical to embryo injections and oocyte handling as described previously by others (Nair S. et al., 2013). Prior to fertilization injected oocytes recovered for 30 minutes at room temperature.

### Statistical Methods

Asterisks indicate the p-values where a Welsh Two-Sample T-test was performed under the assumption of unequal variance and non-paired samples. Default statistical thresholds (p<0.01) were applied as part of DESeq2 to identify differentially expressed genes in RNA-Seq analysis. ChIP-Seq peaks were regions with greater than 3-fold enrichment over input lambda as assigned by Macs2. For correlational analysis a Pearson test was applied on complete datasets, regions were data was missing were excluded, R-value are indicated where applicable. P-values for Gene Ontology analysis were generating using DAVID default settings.

